# Arabidopsis *mlo3* mutant plants exhibit spontaneous callose deposition and signs of early leaf senescence

**DOI:** 10.1101/558122

**Authors:** Stefan Kusch, Susanne Thiery, Anja Reinstädler, Katrin Gruner, Krzysztof Zienkiewicz, Ivo Feussner, Ralph Panstruga

## Abstract

The family of Mildew resistance Locus O (MLO) proteins is best known for its profound effect on the outcome of powdery mildew infections: when the appropriate MLO protein is absent, the plant is fully resistant to otherwise virulent powdery mildew fungi. However, most members of the MLO protein family remain functionally unexplored. Here, we investigate *Arabidopsis thaliana MLO3*, the closest relative of *AtMLO2, AtMLO6* and *AtMLO12*, which are the Arabidopsis *MLO* genes implicated in the powdery mildew interaction. The co-expression network of *AtMLO3* suggests association of the gene with plant defense-related processes such as salicylic acid homeostasis. Our extensive analysis shows that *mlo3* mutants are unaffected regarding their infection phenotype upon challenge with the powdery mildew fungi *Golovinomyces orontii* and *Erysiphe pisi*, the oomycete *Hyaloperonospora arabidopsidis*, and the bacterial pathogen *Pseudomonas syringae* (the latter both in terms of basal and systemic acquired resistance), indicating that the protein does not play a major role in the response to any of these pathogens. However, *mlo3* genotypes display spontaneous callose deposition as well as signs of early senescence in six-or seven-week-old rosette leaves in the absence of any pathogen challenge, a phenotype that is reminiscent of *mlo2* mutant plants. We hypothesize that de-regulated callose deposition in *mlo3* genotypes is the result of a subtle transient aberration of salicylic acid-jasmonic acid homeostasis during development.

## Introduction

Members of the family of Mildew resistance Locus O (MLO) proteins are well known for their intriguing effect regarding the modulation of susceptibility to powdery mildew fungi in plants. Loss-of-function mutants of the appropriate *MLO* gene(s) confer durable broad-spectrum resistance against powdery mildews (1), an effect originally discovered in barley (2) and later validated in many monocotyledonous and dicotyledonous plant species (3).

In Arabidopsis, three genes (*AtMLO2*, At1g11310; *AtMLO6*, At1g61560; and *AtMLO12*, At2g39200) unequally contribute to susceptibility to powdery mildews. The *mlo2* loss-of-function mutation alone causes macroscopically discernable resistance to the powdery mildew species *Golovinomyces cichoracearum* and *G. orontii*, but retains around 30-50 % entry rate by these fungi (4). Both *mlo6* and *mlo12* mutations potentiate the resistance phenotype of *mlo2* mutants, but do not confer enhanced resistance to powdery mildew on their own. Only the triple mutant *mlo2 mlo6 mlo12* is fully resistant, with <1 % entry success by the fungal invader (4). However, at least in barley and Arabidopsis the resistance trait is associated with pleiotropic phenotypes. Both the barley *mlo* and the Arabidopsis *mlo2* mutant exhibit spontaneous deposition of distinct callose aggregations in ageing but otherwise pathogen-free leaves (5), as well as early leaf senescence. The latter is first exemplified by mesophyll cell death (6) and later by catabolism of photosynthetic pigments, leaf chlorosis and necrosis (4, 5, 7, 8). Furthermore, the barley *mlo* and Arabidopsis *mlo2 mlo6 mlo12* triple mutants are altered in the infection phenotype to a range of microbial phytopathogens unrelated to powdery mildews (9, 10). The Arabidopsis *mlo2* single mutant also lacks systemic acquired resistance (SAR) against the bacterial pathogen *Pseudomonas syringae* (11) and exhibits enhanced tolerance to ozone (12). The barley *Mlo* gene codes for a seven-transmembrane domain protein with an extracellular or luminal amino-terminus and a cytosolic carboxy-terminus (13, 14). The carboxyterminal tail contains an evolutionarily conserved calmodulin binding site that is required for full functionality of barley Mlo towards barley powdery mildew infection (15). In addition, four conserved cysteine residues in the first and third extracellular loop are required for barley Mlo function in the context of the powdery mildew interaction (16), and the short tetrapeptide motif [DE]FSF at the carboxy-terminus is a characteristic feature of MLO proteins involved in powdery mildew susceptibility (17). Several additional peptide motifs have been found that are highly conserved throughout MLO proteins of land plants (18, 19).

Comprehensive phylogenetic analyses of the MLO protein family have shown that MLO proteins can be grouped in at least seven distinct clades (19–21). All MLO proteins known to play a role in the powdery mildew interaction belong to clade IV in monocotyledonous plants (e.g. barley Mlo) and clade V in dicotyledonous plants (such as *At*MLO2, *At*MLO6, and *At*MLO12). The Arabidopsis genome encodes 15 MLO family members, represented by five of these clades (22). Apart from the interaction with powdery mildews, Arabidopsis MLO proteins have been implicated in root thigmomorphogenesis as the clade I *mlo* mutants *mlo4* and *mlo11* display a root curling phenotype upon a tactile stimulus *in vitro* (23, 24). The clade III *mlo* mutant *nortia* (*nta, mlo7*) causes reduced fertility since the female gametophyte is impaired in pollen tube recognition, resulting in pollen tube overgrowth in the synergids (25). *At*NTA/*At*MLO7 function requires localization to a Golgi-associated compartment, and the protein forms homo-oligomers during pollen tube perception (26).

*At*MLO3 (At3g45290; clade VI) is closely related to *At*MLO2, *At*MLO6, and *At*MLO12 (19), but a respective *mlo3* knockout mutant did not display increased resistance to *G. orontii* at the macroscopic level (4). A β-glucuronidase (GUS) reporter construct under control of the *AtMLO3* promoter revealed a similar expression pattern as a respective reporter construct driven by the *AtMLO2* promoter, suggesting potentially overlapping or redundant function(s) of the two proteins (22). However, *AtMLO3* and its encoded protein have never been functionally analyzed in detail. In this study, we expanded the analysis of the *AtMLO3* expression profile and identified a set of co-expressed genes that overlap with those co-expressed with *AtMLO2*. We further characterized a set of *mlo3* Transfer-DNA (T-DNA) insertion mutants with regard to the interaction with phytopathogens from various kingdoms of life. These comprised two powdery mildew fungi (the adapted species *G. orontii* and the non-adapted pea powdery mildew pathogen *Erysiphe pisi*), the oomycete *Hyaloperonospora arabidopsidis* (*Hpa*), and the bacterium *P. syringae* pv. *tomato* (*Pst*). We further tested if, like in the case of *mlo2, mlo3* mutants are impaired in SAR to *P. syringae* pv. *maculicola* (*Psm*). In addition, we analyzed whether *mlo3* mutants, similar to *mlo2* mutants, exhibit spontaneous callose deposition and early leaf senescence in unstressed conditions. Our data revealed no apparent role for *AtMLO3* in plant immunity, but the findings suggest an overlapping function with *AtMLO2* regarding the control of callose deposition and the appropriate onset of leaf senescence.

## Results

### Arabidopsis *MLO3* is co-expressed with defense response genes

*AtMLO3* is phylogenetically the closest relative of *AtMLO2, AtMLO6*, and *AtMLO12* (19), and according to promoter-GUS studies the four genes seem to be largely co-expressed (22). We used the electronic fluorescent pictograph (eFP) browser online tool (27) for a more detailed analysis of the *AtMLO3* expression pattern. We noted high *AtMLO3* transcript levels in the senescing leaf (Figure S1) and in the context of various biotic stresses. These comprise sites of *G. orontii* infection at 5 days post infection (dpi), challenge with the incompatible pathogens *Pst* DC3000 Δ*hrcC* and *Hpa* Emwa2 (in the Arabidopsis *rpp4* mutant, which is susceptible to Emwa2), and treatment with the microbe-associated molecular patterns (MAMPs) flg22 and HrpZ. Weak *AtMLO3* expression was further seen in the shoot during osmotic stress (Figure 1A). In addition, both *AtMLO2*, which was shown to exhibit a critical role during SAR (11), and *AtMLO3* were upregulated in systemic leaves upon primary *Psm* inoculation, as well as after irrigation with pipecolic acid in a *FLAVIN MONOOXYGENASE1* (*AtFMO1*)-dependent manner (Table S1; data compiled from (31, 32)). Pipecolic acid is a competent inducer of a SAR-like response and defense priming (31, 33). Together, the expression profile suggests an association of *AtMLO3* function with leaf senescence, pathogen defense -in particular SAR, and/or possibly osmotic stress.

**Fig 1.**
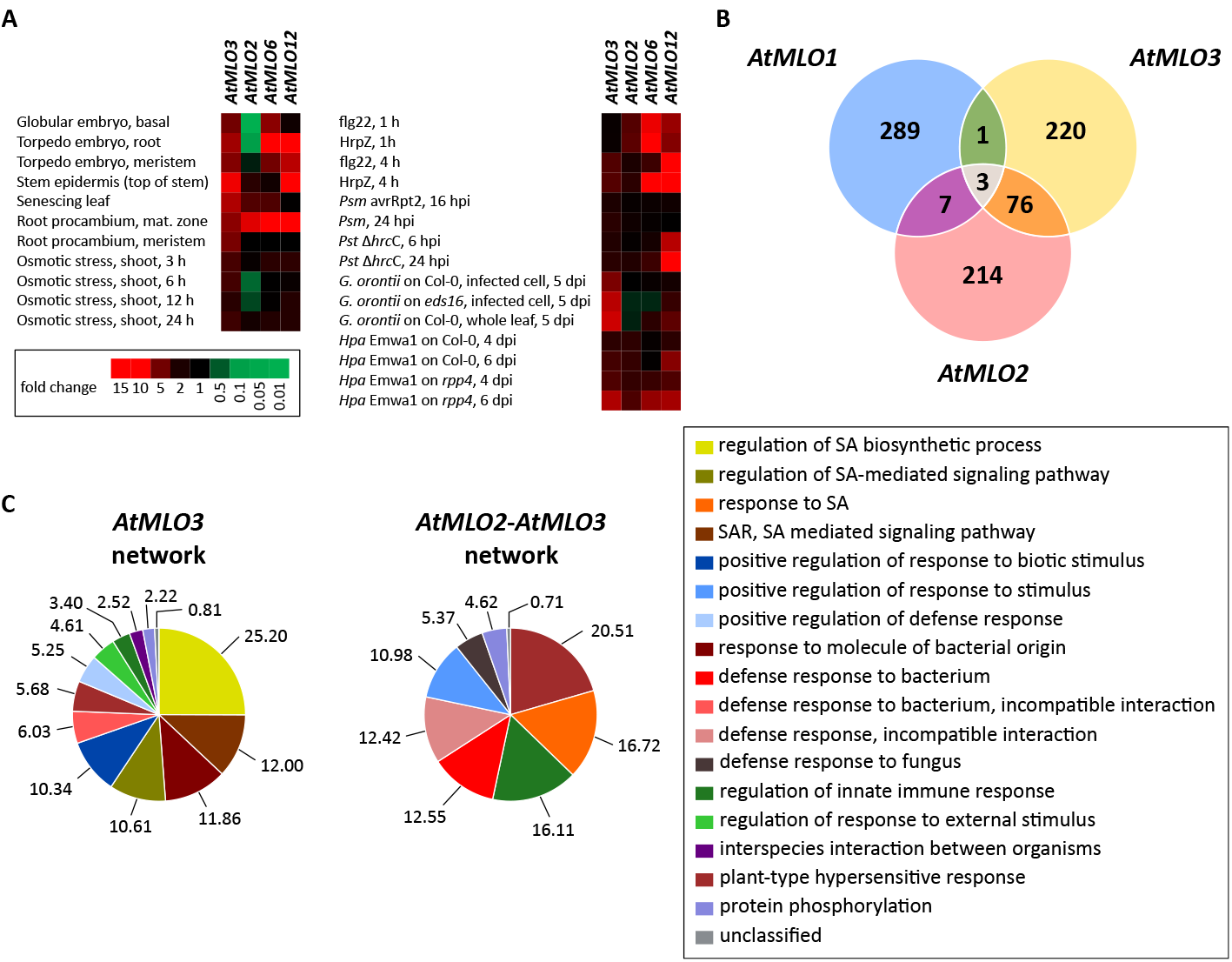
*AtMLO3* is co-expressed with components of plant immunity. **A** Heatmap summarizing the relative expression in fold change of *AtMLO2* (At1g11310), *AtMLO3* (At3g45290), *AtMLO6* (At1g61560), and *AtMLO12* (At2g39200) upon the respective condition. Black, no change; red, increased expression level; green, decreased expression level. Data was extracted using the eFP browser (27). **B.** The top 300 co-expressed genes of *AtMLO1* (At4g02600), *AtMLO2*, and *AtMLO3* were extracted using ATTED-II release 2017.12.14 (28). The Venn diagram summarizes the number of overlapping genes. **C.** Venn diagrams highlighting enriched GeneOntology (GO) terms among the top 300 co-expressed genes of *AtMLO3* (left) and among the 79 genes co-expressed in both *AtMLO2* and *AtMLO3* (middle). The legend to the right indicates the colors that are assigned to each term; the numbers indicate the percentage of genes in the respective network with that term. Gene enrichment analysis was done with PLAZA v4.0 (29) and GO terms were extracted using PANTHER v10 (30).

Next, we extracted the respective co-expression networks for *AtMLO2, AtMLO3*, and *AtMLO1* (clade II *MLO*, which served as a negative control) using the ATTED-II online tool (28). Based on these data (Supplementary File 1) we performed gene ontology (GO) enrichment analysis with PLAZA 4.0 (29). The co-expression networks of *AtMLO2* and *AtMLO3* overlap by 26 %, i.e. 79 out of the 300 top co-expressed genes (Figure 1B). By contrast, there was little overlap between the co-expressed genes of *AtMLO2* and *AtMLO3*, respectively, and those co-expressed with *AtMLO1* (only 11 genes in total). Using PANTHER (release 2018_04; (30)), which provides a hierarchical GO term output, we analyzed the co-expression networks of *AtMLO2, AtMLO3*, and the 79 overlapping genes for over-representation of GO terms (Figure 1C, Table 1 and Supplementary File 1). Terms attributed to biotic stress response, incompatible interaction with pathogens, salicylic acid (SA) response, and SAR were highly represented in the co-expression networks of both genes. In case of *AtMLO3* analyzed alone, SA biosynthesis and regulation were highly over-represented as well. Genes co-expressed with both *AtMLO2* and *AtMLO3* encode a number of proteins with well-defined functions in plant immunity (Supplementary File 1). For example, 20 genes (25 %) harbor the GO term “response to biotic stress”, 27 genes (34 %) are annotated with “defense response”, and eight genes are associated with “response to salicylic acid” (Supplementary File 2). The shared set comprises genes encoding a range of receptor-like kinases, including prominent members such as *At*BIR1 and *At*SOBIR1 (34), components of SA signaling (*At*EDS1; (35)), transcriptional regulators of SA responses (*At*NPR4; (36, 37)), NBS-LRRs including *At*RPS2 (38, 39), as well as some WRKY transcriptional regulators with known roles in plant defense and/or as negative regulators of plant senescence (*At*WRKY53, *At*WRKY54 and *At*WRKY70; (40–42)).

**Table 1.** Highly over-represented biological process GO terms in the common co-expression network of *AtMLO2* and *AtMLO3*.

| GO-term | Log2-Enrichment | p-value | Description |
| --- | --- | --- | --- |
| GO:1902478 | 6.31 | 8.21E-04 | Negative regulation of defense response to bacterium, incompatible interaction |
| GO:0080142 | 6.16 | 2.80E-05 | Regulation of salicylic acid biosynthetic process |
| GO:0009697 | 5.67 | 8.04E-05 | Salicylic acid biosynthetic process |
| GO:0009862 | 5.31 | 1.74E-04 | Systemic acquired resistance, salicylic acid mediated signaling pathway |
| GO:0010337 | 5.21 | 2.16E-04 | Regulation of salicylic acid metabolic process |
| GO:0031098 | 5.21 | 2.16E-04 | Stress-activated protein kinase signaling cascade |
| GO:0010112 | 5.11 | 2.65E-04 | Regulation of systemic acquired resistance |
| GO:0046189 | 4.82 | 4.87E-04 | Phenol-containing compound biosynthetic process |
| GO:0042391 | 4.78 | 5.26E-04 | Regulation of membrane potential |
| GO:0009682 | 4.67 | 6.56E-04 | Induced systemic resistance |
| GO:0009696 | 4.51 | 9.14E-04 | Salicylic acid metabolic process |
| GO:0048544 | 4.35 | 1.32E-04 | Recognition of pollen |
| GO:0009816 | 4.32 | 1.66E-06 | Defense response to bacterium, incompatible interaction |
| GO:0008037 | 4.27 | 1.63E-04 | Cell recognition |
| GO:0031348 | 4.26 | 1.88E-05 | Negative regulation of defense response |
| GO:0009626 | 4.22 | 2.53E-06 | Plant-type hypersensitive response |
| GO:0034050 | 4.21 | 2.66E-06 | Host programmed cell death induced by symbiont |
| GO:0009875 | 4.21 | 1.91E-04 | Pollen-pistil interaction |
| GO:0009627 | 4.17 | 2.54E-05 | Systemic acquired resistance |

We then assessed the 1000 base pair (bp) upstream regions of the 79 co-expressed genes from the common co-expression network of *AtMLO2* and *AtMLO3* for known and/or common *cis*-regulatory elements. Using the GO annotations from PLAZA4.0 (Supplementary File 2), we further subdivided these genes into subgroups connected to defense or immune response, SAR, and SA-related processes. On the basis of analysis with AthaMap (43, 44), which performs database-assisted searches for known *cis*-regulatory elements, we found that *At*WRKY6 (five genes) and *At*WRKY18 (seven genes) binding motifs were more than 4-fold over-represented in the set of 79 genes (Supplementary File 3). Among the defense-related genes, *At*WRKY6 was only found once, while other putative WRKY binding sites (*At*WRKY12, *At*WRKY18, *At*WRKY38, and *At*WRKY43) as well as two MYB factor binding sites (*At*MYB.PH3, *At*MYB98) occurred multiple times each. Next, we subjected the 1000 bp upstream sequences of the 79 genes to an unbiased detection of sequence motifs using MEME v5.0.4 (45). While the query sequences derived from genes associated with the GO terms SA and SAR did not yield any significant motifs at p < 1e-2, two non-homopolymeric motifs were overrepresented in the upstream regions of genes sharing GO terms related to defense response (Supplementary File 4). Using the MEME-TOMTOM tool against the *Arabidopsis thaliana* DAP motif database, we found that these may represent *At*WRKY47 and *At*RAP2.12 binding sites. Overall, our *cis*-regulatory element analysis suggests that the co-expression network of *AtMLO2* and *AtMLO3* might be mainly regulated by WRKY transcription factors, in particular those genes in this network related to the plant defense response.

We next used a transgenic promoter-GUS (P*AtMLO3*::*GUS*) line, and as a control a P*AtMLO2*::*GUS* line (22), to examine whether the expression pattern revealed by *in silico* analysis can be detected histologically upon ageing, pathogen stress, wounding, or MAMP treatment. As previously reported (7), strong GUS signals were seen in ageing rosette leaves of the P*AtMLO2*::*GUS* line; however, we found hardly any detectable GUS signal with the P*AtMLO3*::*GUS* line, not even in eight-week-old plants grown under long day conditions (Figure S2A). In case of the *AtMLO3*::*GUS* line, GUS staining was detectable in spots in old rosette leaves of plants grown under short day conditions, in trichomes, and occasionally in the meristematic zone of small leaves (Figure S2B). Similarly, only the P*AtMLO2*::*GUS* line was responsive to wounding by tweezers, with GUS signal detectable as late as 24 h after wounding, while the P*AtMLO3*::*GUS* plants showed no recognizable response (Figure S2C). To test the reaction of the two GUS reporter lines to MAMPs, we treated seedlings grown in liquid Murashige and Skoog (MS) medium with flg22 (100 nM) or chitin (100 µg mL^-1^) and found the P*AtMLO2*::*GUS* line to be somewhat responsive, while we observed no GUS signal in P*AtMLO3*::*GUS* seedlings (Figure S2D). Both transgenic promoter-GUS lines lacked a detectable increase in GUS staining upon challenge with virulent pathogens (*G. orontii, Hpa*, or *Pst* DC3000), equally upon macroscopic inspection and following microscopic examination of infection sites (Figure S2E). Taken together, publicly available microarray and co-expression data suggests that *AtMLO3* might be associated with senescence and biotic stress responses, while the transgenic promoter-*GUS* line did not support this notion.

### Mutations in *AtMLO3* do not affect compatible and incompatible powdery mildew interactions

Since *At*MLO3 is closely related to clade V MLO proteins like *At*MLO2, *At*MLO6, and *At*MLO12, we hypothesized that *At*MLO3 might also be involved in the modulation of powdery mildew interactions. We used four independent T-DNA insertion lines as single mutants (*mlo3*-2 (4), *mlo3*-4, *mlo3*-5 and *mlo3*-6) to test this possibility experimentally (Figure 2A). We failed to detect the full-length transcript of *AtMLO3* in these lines by reverse transcription-polymerase chain reaction (RT-PCR) analysis, indicating that they are true knockout mutants (Figure 2B). Furthermore, we created double and higher order mutants of *mlo3*-4 in combination with *mlo2*-5, *mlo6*-2, and *mlo12*-1. When challenged with the adapted powdery mildew pathogen *G. orontii*, all four *mlo3* single mutant lines showed similar susceptibility as the wild type accession Col-0, with about 80 % entry success at 48 hours post inoculation (hpi) and intense sporulation at 7 dpi (Figure 3A and Figure S3). Similarly, the combinations of *mlo3*-4 with *mlo2*-5, *mlo6*-2, and *mlo12*-1 exhibited the same levels of penetration success at 48 hpi as the respective mutant line with wild type *MLO3* present (Figure 3B), and all plants harboring the *mlo2*-5 mutation were without macroscopically visible powdery mildew symptoms at 7 dpi (Figure S3). Next, we tested the interaction of *mlo3* mutants with the non-adapted pea powdery mildew pathogen *Erysiphe pisi*. Upon challenge with this fungus, the *mlo3* single mutants displayed similar entry rates of about 15 % as the Col-0 wild type. The combination with *mlo2*-5 did not change the entry rate significantly (Figure 3C), suggesting that *AtMLO3* also does not play a prominent role in defense against the non-adapted powdery mildew pathogen.

**Fig 2.**
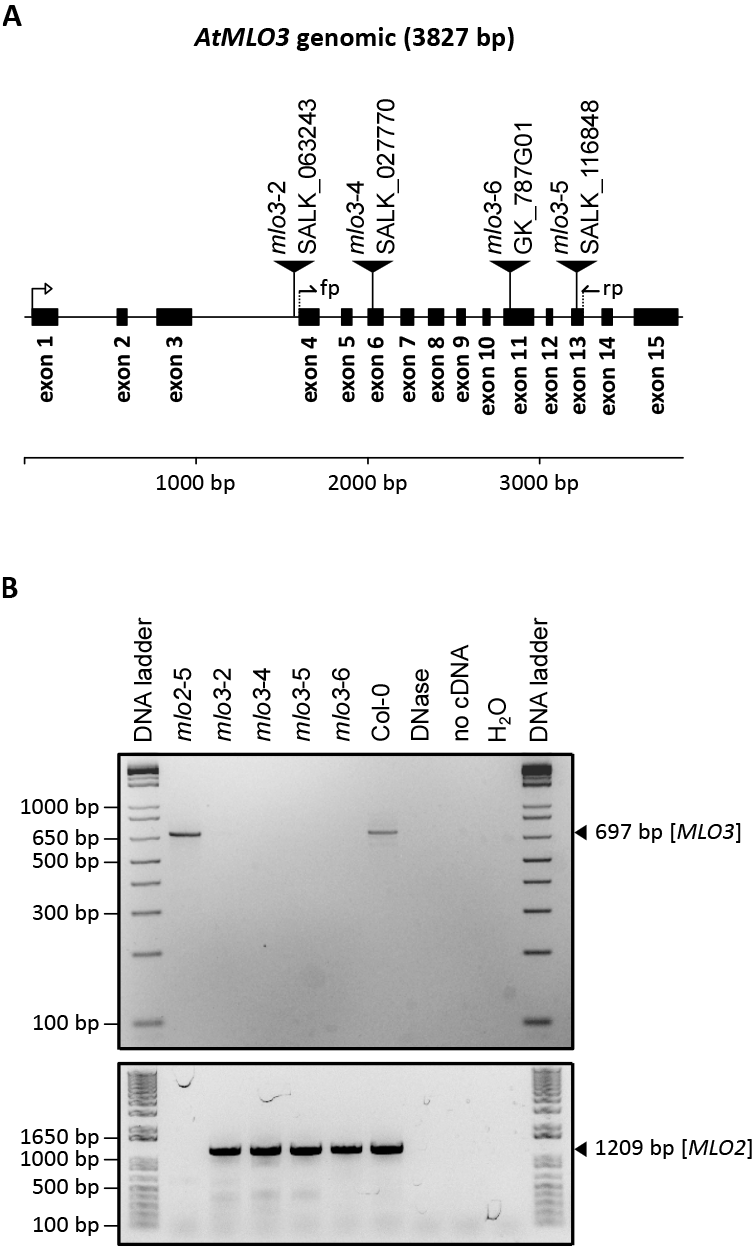
Arabidopsis *mlo3* T-DNA insertion lines do not express *AtMLO3*. A. **A** Genomic map of *AtMLO3*. Black boxes represent exons, the lines in between introns. The location of the respective T-DNA insertions is indicated by triangles. fp, forward primer (AtMLO_3C); rp, reverse primer (AtMLO_331). **B.** Upper panel, agarose gel for the RT PCR of the *AtMLO3* transcript. The band is expected at 697 bp (fp + rp), as indicated on the right. Lower panel, agarose gel for the RT-PCR of the *AtMLO2* transcript (as control). The size of the band is 1,209 bp, as indicated on the right. Primers used are as described in (9). On the left, spacing of standard DNA fragments is indicated. DNA ladder, 1 Kb Plus DNA ladder (Invitrogen-Thermo Fisher, Waltham, MA, USA); DNase, water instead of RNA used for DNase digestion as template (negative control); no cDNA, water instead of RNA used for cDNA synthesis (negative control); H2O, water template (negative control).

**Fig 3.**
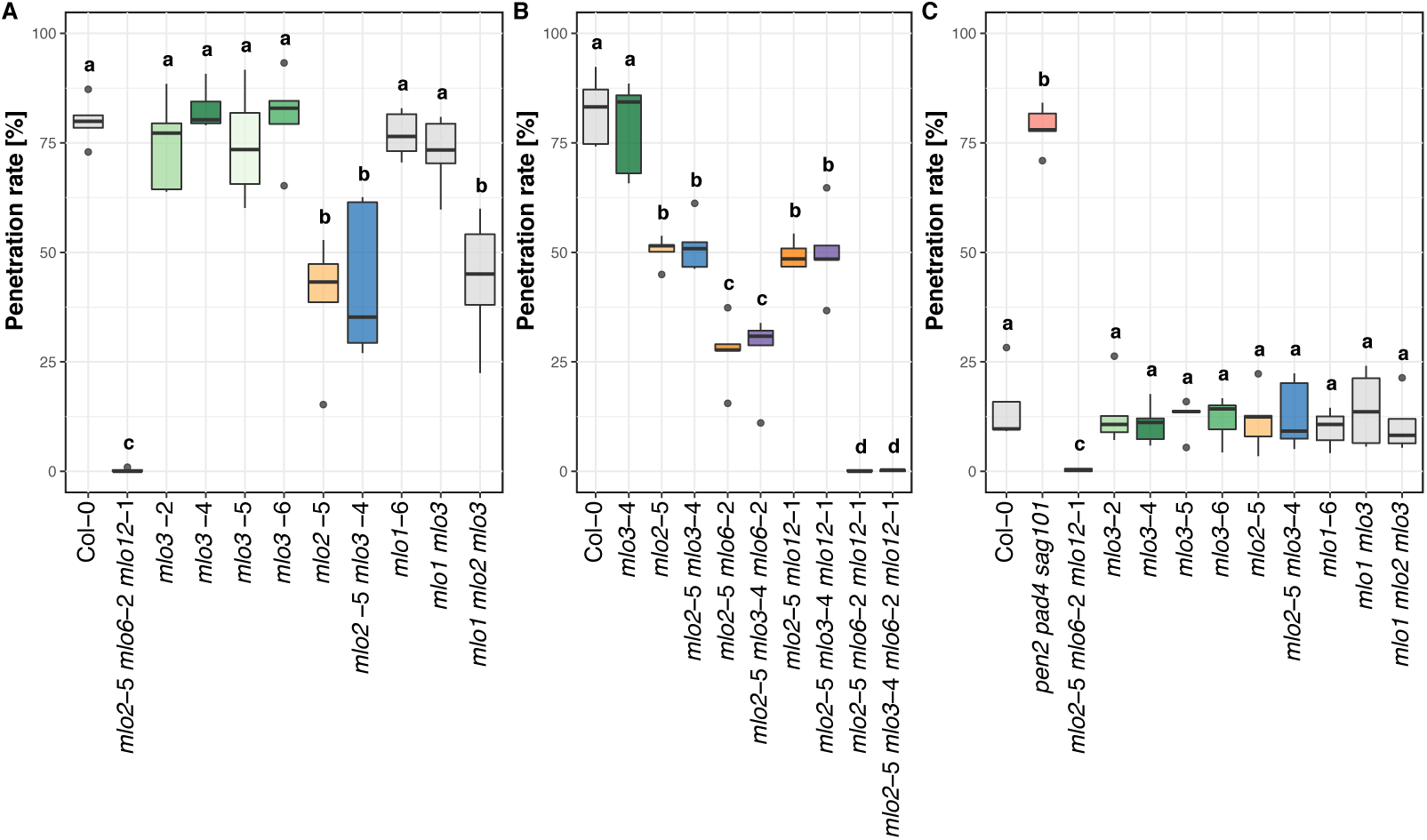
Arabidopsis *mlo3* mutants show unaltered powdery mildew infection phenotypes. Plants were inoculated with the indicated powdery mildew pathogen and the penetration rate was determined at 48 hpi. **A.** Boxplot showing *G. orontii* penetration rates for the *mlo3* mutant lines (Figure 2A) and for mutant combinations with *mlo2*-5 and *mlo1*-6. Col-0 is the wild type, *mlo2*-5 *mlo6*-2 *mlo12*-1 the resistant control, *mlo2*-5 and *mlo1*-6 are parental controls for the mutant combinations. **B.** Boxplot showing penetration success of *G. orontii* for mutant combinations of *mlo3*-4 with *mlo2*-5, *mlo6*-2 and *mlo12*-1. Col-0 and *mlo3*-4 are susceptible controls, the respective parent control harboring wild type *AtMLO3* is always placed left from the *mlo3*-4-containing mutant combination. **C.** Boxplot for the *E. pisi* penetration rate with the same mutants as in (**A**), including the hyper-susceptible control *pen2*-1 *pad4*-1 *sag101*-1 (46). Each replicate is based on 150-200 interaction sites scored on four to five leaves from one leaf per plant and genotype; data shown is based on the means from each replicate. n = 5 independent replicates in all experiments. Statistical testing was done with GLM (binomial distribution).

We further performed transient overexpression of *At*MLO2 C-terminally tagged with yellow fluorescent protein (*At*MLO2-YFP) and *At*MLO3-YFP in Arabidopsis leaf epidermal cells to analyze comparatively the subcellular localization of the two *At*MLO proteins. *At*MLO2-YFP localized to the cell periphery and was additionally present in discrete vesicular structures in the cells, while *At*MLO3-YFP seemed to reside exclusively at the cell periphery (Figure 4). We found that *At*MLO2-YFP re-localized to and accumulated at penetration sites of *G. orontii*, while subcellular localization of *At*MLO3-YFP was seemingly unaffected by the challenge with this pathogen (Figure 4). Altogether, our data indicates that *AtMLO3* is not involved in the interaction between Arabidopsis and powdery mildew fungi.

**Fig 4.**
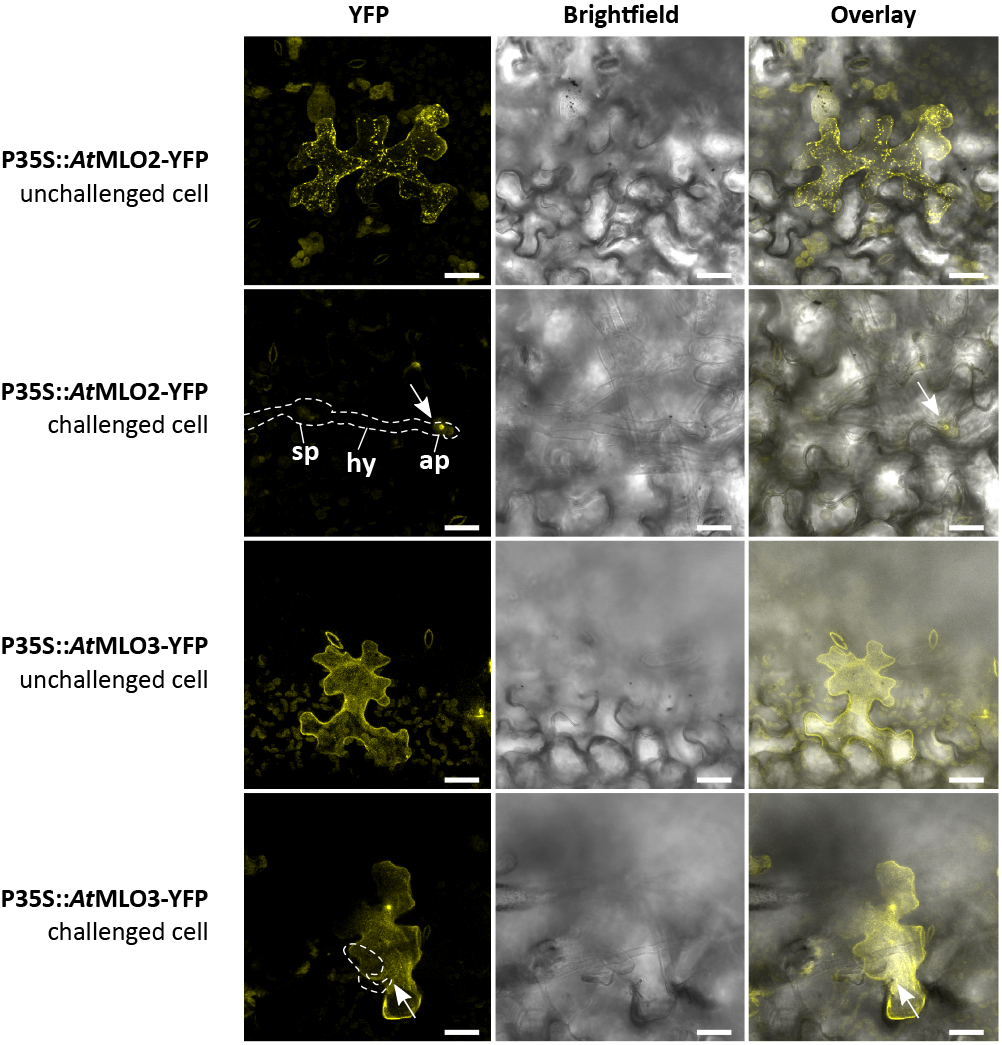
*At* MLO3-YFP does not re-localize to powdery mildew penetration sites. *AtMLO2*-YFP (upper two panels) and *AtMLO3*-YFP (lower two panels) were transiently over-expressed under control of the cauliflower mosaic virus 35S promoter in leaf epidermal cells of Arabidopsis *mlo2*-5 *mlo3*-4 *mlo6*-2 *mlo12*-1 plants by use of biolistic gold particle delivery. “Challenged cell” designates cells challenged with *G. orontii* at 48 hpi (fungal structures indicated by the dashed line). The left micrographs show the YFP signal (YFP excitation at 514 nm, emission at 525-570 nm), the middle micrographs represent the respective brightfield image and the right micrographs depict the overlay of both. Confocal YFP images were generated by 3D reconstruction and are shown as maximum projection of all layers. Arrows point at the *G. orontii* penetration sites. sp, spore; ap, appressorium; hy, secondary hyphae. Scale bar: 25 µm.

### *mlo3* mutants show wild type-like colonization by *Hpa* and *Pst*

Due to the apparent association of *AtMLO3* with components and processes of plant defense (Figure 1), we suspected that despite the fact that powdery mildew susceptibility was unaffected in *mlo3* mutant plants, *AtMLO3* might play a role in modulating defense to other pathogens. To test this idea, we employed the hemi-biotrophic bacterial pathogen *Pst* DC3000 and the biotrophic oomycete *Hpa* Noco2 in our experiments. *Hpa* Noco2 (virulent on Col-0) was spray-inoculated and the reproductive success was determined as (conidio-)spores per g fresh weight at 7 dpi. We found that all tested genotypes (*mlo3* single mutants, *mlo1*-6, *mlo2*-5, and combinations of *mlo3*-4 with *mlo1*-6 and/or *mlo2*-5) allowed oomycete sporulation at levels of 2.5-7.5 x 10^5^ spores/g FW (Figure 5A), which is comparable to the Col-0 wild type. The Arabidopsis accession Landsberg *erecta* (L-*er*; harboring the *Hpa* resistance gene *RPP5*) was fully resistant and the *eds1*-2 [Col-0] mutant was more susceptible (*ca.* 10 x 10^5^spores/g FW) to *Hpa* Noco2, as previously reported for these genotypes (47, 51). The result of this experiment indicates that neither *mlo2* nor *mlo3* mutants show an altered infection phenotype to the adapted oomycete *Hpa* Noco2.

**Fig 5.**
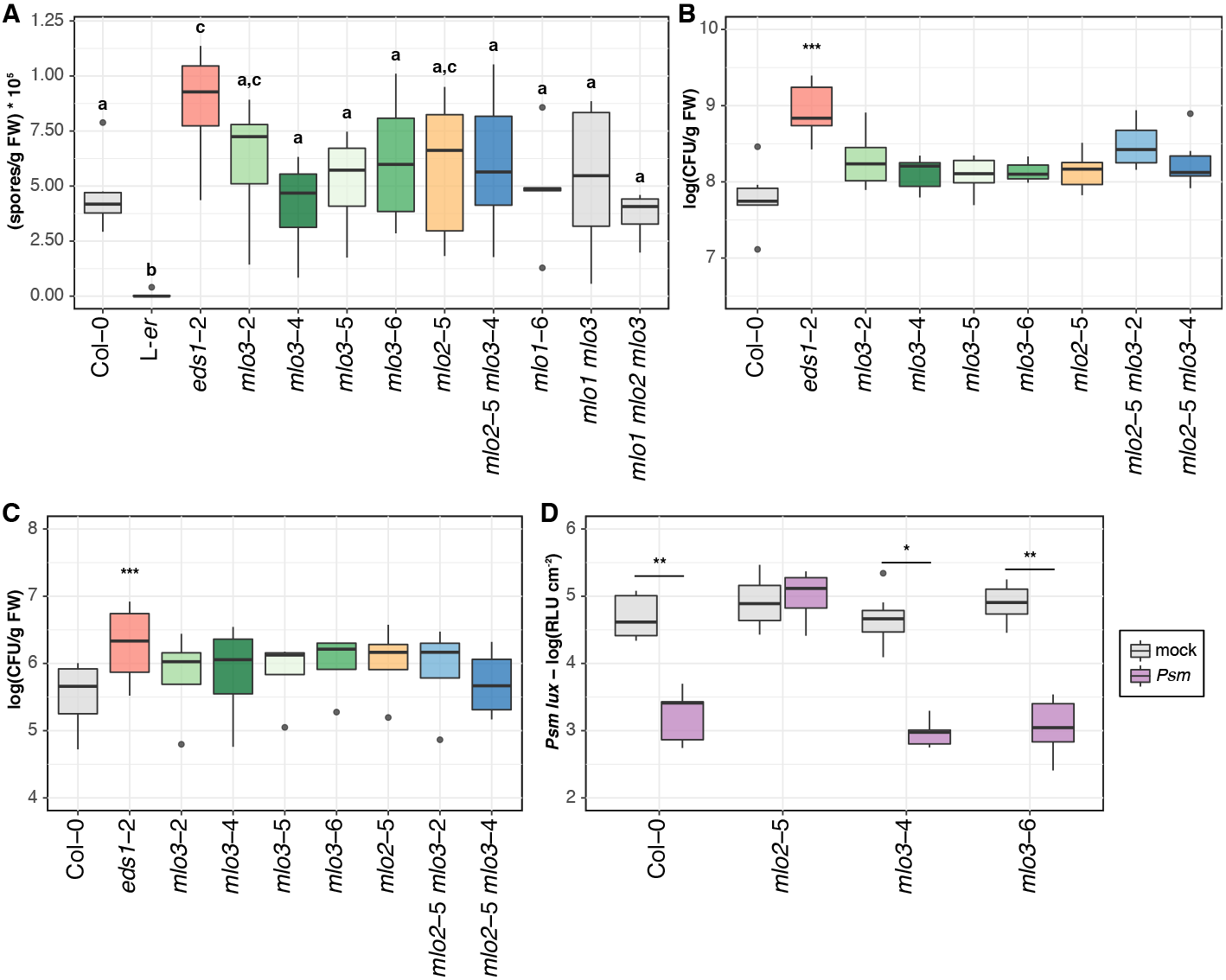
Arabidopsis *mlo3* mutants show unaltered *Hpa* and *Pst* infection phenotypes and are SAR-competent in association with *Psm*. **A** *Hpa* sporulation measured at 7 dpi in (conido-)spores per gram fresh weight (FW). Genotypes are as in Figure 3A, with Col-0 being the susceptible wild type, accession L-*er* the resistant control (47), and *eds1*-2 in Col-0 background (48) the hyper-susceptible control. n = 6 independent replicates, and GLM (Poisson distribution) was used for statistical testing. **B.** *Pst* DC3000 bacterial proliferation was determined at 3 dpi in Arabidopsis seedlings after spray inoculation. The bacterial titer is expressed as log10 of colony-forming units (CFU) per gram fresh weight. Col-0 is the wild type, *mlo2*-5 a parental and *eds1*-2 a hyper-susceptible control. The four *mlo3* mutant lines and the double mutants *mlo2*-5 *mlo3*-2 and *mlo2*-5 *mlo3*-4 were subjected to *Pst* inoculation. n = 6 independent replicates with GLM (Poisson distribution) as statistical test. **C.** Data of bacterial infection assay as in (**B**), with the avirulent strain *Pst* DC3000 Δ*hrcC* (49, 50). n = 4 independent replicates using GLM (Poisson distribution) as statistical test. **D.** SAR was monitored by primary infiltrations of three lower leaves of five-week-old plants with *Psm* or 10 mM MgCl_2_ (mock), followed by inoculations of systemic leaves with *Psm lux* two days later. Bacterial proliferation of *Psm lux* was measured at 3 dpi, expressed as relative light units (RLU) cm^-2^. n = 1 biological replicate with nine plants per genotype, statistical testing was performed by GLM (quasi-Poisson distribution model). Similar results were obtained in a second independent experiment (Figure S4).

*Pst* DC3000 (virulent) and its derived mutant strain *Pst* DC3000 Δ*hrcC* (lacking the type III secretion apparatus and being strongly compromised in virulence; (50)) were inoculated on 16-day-old Arabidopsis seedlings by spray-inoculation, and the bacterial titer in rosette leaves was determined as colony-forming units (CFU) at 0 dpi (inoculation control; see Figure S4) and at 3 dpi. The Col-0 wild type displayed a bacterial titer of *ca.* 10^8^CFU/g FW, while the hyper-susceptible control *eds1*-2 [Col-0] (48) had a significantly enhanced titer of around 10^9^CFU/g FW (Figure 5B). The *mlo3* and *mlo2*-5 single mutants as well as the *mlo2*-5 *mlo3*-2 and *mlo2*-5 *mlo3*-4 double mutants all showed a slightly increased bacterial titer compared to Col-0 (>10^8^CFU/g FW), but the observed differences were not significant in our set of experiments according to thorough statistical testing. Similarly, all these mutant lines revealed bacterial titers comparable to Col-0 following inoculation with *Pst* DC3000 6.*hrcC* (around 10^6^CFU/g FW); only *eds1*-2 [Col-0] was more susceptible with *ca.* 10^7^CFU/g FW (Figure 5C). Finally, since both publicly available microarray co-expression data (Table 1) and deep transcriptome sequencing (RNA-Seq) data (Table S1) indicate a possible functional association of *AtMLO3* with SAR, we investigated the ability of *mlo3* mutant lines to exhibit SAR against *Psm*. Three primary leaves of five-week-old plants were pressure-infiltrated with the virulent *Psm* strain ES4326, followed by inoculation of three systemic leaves with *Psm lux* (a transgenic variant of *Psm* constitutively expressing luciferase; (52)) two days later. Bacterial proliferation of *Psm lux* was quantified at 3 dpi by measuring luciferase activity (given as relative light units, RLU). Both *mlo3* mutant lines were competent in acquiring resistance to *Psm lux* after pre-inoculation with *Psm* in a wild type-like manner, with approximately 10^3^RLU cm^-2^after preinfection with *Psm*, compared to *ca.* 10^5^RLU cm^-2^after mock treatment (Figure 5D and Figure S4C). By contrast, the *mlo2*-5 mutant revealed similar RLU values upon both mock and *Psm* pre-treatment, since this mutant is SAR-incompetent (11). Together, these data suggest that Arabidopsis *mlo3* mutants are not altered regarding colonization by the bacterial pathogen *Pst* DC3000 and they do not show a lack of SAR against *Psm lux* in our experimental conditions.

### *mlo3* mutants are not affected in development upon osmotic stress

Guided by the slightly induced expression levels of *AtMLO3* in roots upon osmotic stress (Figure 1A), we determined the osmotic stress tolerance of *mlo3* mutant lines *in vitro*. Seedlings grown on MS plates containing 300 mM of mannose were quantitatively assessed for root length, the number of lateral roots, fresh weight, and leaf number (developmental stage). We found that with a few outliers, the *mlo2* and *mlo3* single mutants as well as the tested *mlo2 mlo3* double mutants behaved like the wild type Col-0 in all respects. The root length of the two-week-old seedlings was in the range of 10-15 mm in our setting, as opposed to *ca.* 50 mm in non-stress conditions. The number of lateral roots was between one and two, the fresh weight was at 1-2 mg per plant, and the leaf number, which was usually four when unstressed, was at least eight (Figure S5). Accordingly, Arabidopsis *mlo3* single mutants do not show a significantly altered development upon exposure to osmotic stress by mannose.

### *mlo3* mutants exhibit spontaneous callose deposition similar to *mlo2*

In addition to enhanced powdery mildew resistance, Arabidopsis *mlo2* mutant plants exhibit an early leaf senescence phenotype, which is associated with spontaneous callose deposition in rosette leaves (4, 7). The early senescence phenotype cannot be seen in *mlo6* and *mlo12* single mutants (4). We analyzed six-week-old *mlo3* mutant plants grown under short day conditions for the development of spontaneous callose depositions in rosette leaves. This was done in comparison to Col-0 wild type and the *mlo2*-5 mutant, which served as negative and positive controls, respectively, in this set of experiments. To this end, we collected all rosette leaves from each individual plant, stained them with aniline blue, photographed randomly selected leaf spots for aniline blue-derived fluorescence, and quantified the number of deposits mm^-2^in the mesophyll (Figure 6A and Figure 6B). We found very few callose deposits in Col-0 (less than 1 mm^-2^), while the *mlo2*-5 mutant showed elevated numbers of these deposits (2-8 mm^-2^). The mutant lines *mlo3*-2 and *mlo3*-4 displayed slightly lower numbers (2-5 mm^-2^), which were nonetheless significantly increased compared to the wild type in five out of six independent experiments. In a separate experiment, comprising also double and higher order mutants, the average callose deposits mm^-2^were somewhat higher for all tested genotypes, with ∼6 (Col-0), ∼12 (*mlo3*-2) and ∼26 (*mlo3*-4). The *mlo2 mlo3* double mutants had on average ∼33 and ∼70 deposits mm^-2^, the *mlo2 mlo6 mlo12* showed ∼99 deposits mm^-2^, and the quadruple mutant *mlo2 mlo3 mlo6 mlo12* displayed ∼55 deposits mm^-2^in rosette leaves of six-week-old plants (Table 2, Figure 6B and Figure S6). We further crossed *mlo3*-2 and *mlo3*-4 with *pmr4*-1, which is deficient in a callose synthase (GLUCAN-SYNTHASE-LIKE5/POWDERY MILDEW RESISTANT4, *At*GSL5/*At*PMR4) that is needed for the deposition of wound and papillary callose (54, 55). *At*PMR4 is further required for the spontaneous callose deposition phenotype of *mlo2* mutant plants (4). Resulting *mlo3 pmr4* double mutants showed wild type-like callose deposition in leaves of six-week-old plants, with *ca.* 2 and 4 deposits mm^-2^, respectively (Table 2 and Figure S6).

**Table 2.** Average and maximum callose deposition in rosette leaves of six-week-old plants.

| Genotype | Replicate 1 |  | Replicate 2 |  |
| --- | --- | --- | --- | --- |
| | Mean ± SD | Maximum | Mean ± SD | Maximum |
| Col-0 | 6.3 (± 3.8) | 16.3 | 3.4 (± 1.7) | 7.6 |
| *mlo3-2* | 25.6 (± 46.2) | 266.3 | 6.4 (± 7.5) | 37.4 |
| *mlo3-2 pmr4-1* | 1.7 (± 2.4) | 9.4 | 2.1 (± 1.9) | 8.1 |
| *mlo3-4* | 11.9 (± 12.4) | 48.1 | 8.0 (± 9.0) | 63.0 |
| *mlo3-4 pmr4-1* | 3.7 (± 4.1) | 16.0 | 2.7 (± 1.9) | 8.5 |
| *mlo2-5* | 59.0 (± 48.6) | 224.3 | 12.8 (± 15.5) | 78.9 |
| *mlo2-5 mlo3-2* | 33.4 (± 38.4) | 135.3 | 8.2 (± 12.3) | 83.2 |
| *mlo2-5 mlo3-4* | 69.2 (± 53.0) | 206.4 | 40.7 (± 41.5) | 174.1 |
| *mlo2 mlo6 mlo12* | 99.8 (± 87.5) | 297.2 | 11.7 (± 18.0) | 85.9 |
| *mlo2 mlo3 mlo6 mlo12* | 55.0 (± 52.8) | 177.1 | 43.5 (± 62.8) | 265.0 |

**Fig 6.**
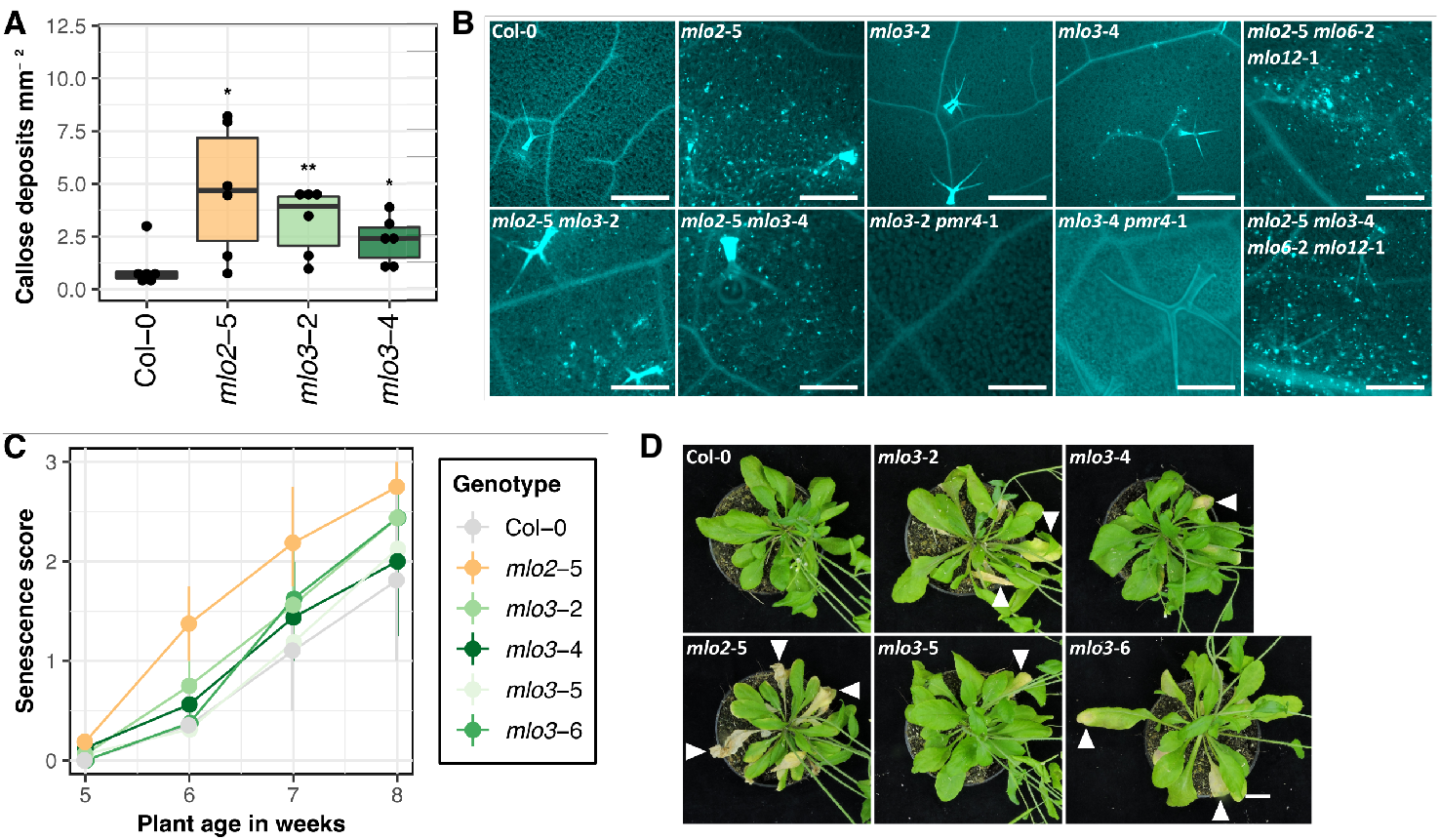
Arabidopsis *mlo3* exhibits spontaneous callose deposition and early leaf senescence in unchallenged rosette leaves. All leaves from six-week-old rosettes were collected and callose was stained using aniline blue. Col-0 wild type served as negative control, while the *mlo2*-5 single and *mlo2*-5 *mlo6*-2 *mlo12*-1 triple mutants were included as positive controls. We tested *mlo3*-2, *mlo3*-4, the respective double mutants with *mlo2*-5, *mlo3*-2 *pmr4*-1, *mlo3*-4 *pmr4*-1, and the quadruple mutant *mlo2*-5 *mlo3*-4 *mlo6*-2 *mlo12*-1. For quantitative assessment, pictures were taken with high black balance and long exposure time to minimize background, and callose deposits were quantified using CellProfiler v3.1.8 (53). **A.** The boxplot shows the average counts for the Col-0 wild type, *mlo3*-2, *mlo3*-4, and *mlo2*-5 from n = 6 independent replicates. Statistics were done with GLM (Poisson distribution). **B.** Representative micrographs for each genotype from (**A**) and Table 2); Scale bar: 500 µm. **C.** The senescence of each plant was rated as follows: 0, no symptoms; 1, mild yellowing symptoms on one or several leaves; 2, pronounced yellowing symptoms on several leaves and/or occasional necrosis; 3, most or all rosette leaves are necrotic. Three to four plants per genotype and replicate were assessed at five, six, seven, and eight weeks after germination; the graph was produced with the average values for each time point and replicate. The error bars indicate maximum and minimum average values. **D.** Representative photographs of seven-week-old rosettes of Col-0, *mlo2*-5, and the *mlo3* mutants. Arrows indicate leaves with visible symptoms of senescence. Scale bar: 1 cm.

Next, we assessed the *mlo3* mutants in comparison to the wild type Col-0 and the *mlo2*-5 mutant for macroscopic signs of early leaf senescence under stress-free long day conditions. Col-0 did not show any visible indication of senescence up to seven weeks after germination, while the *mlo2*-5 mutant displayed extensive senescence symptoms (leaf chlorosis and necrosis) from six weeks onwards (Figure 6C and Figure 6D). We found pronounced senescence symptoms in all four *mlo3* mutants at seven weeks of age, although apparently less drastic than in case of the *mlo2* genotype. Altogether, the data shows that Arabidopsis *mlo3* mutants, like *mlo2* genotypes, exhibit spontaneous and *At*PMR4-dependent callose deposition and early signs of senescence in unchallenged leaves of mature rosettes.

### Levels of SA(G) over JA-IIe are transiently increased in the leaves of *mlo3* mutant plants

Enhanced SA and SA glucoside (SAG) levels were previously observed in *mlo2* mutant plants (11, 56), and SA homeostasis processes were among the highly represented GO terms in the co-expression network of *AtMLO3* (Figure 1C). We therefore measured levels of the phytohormones SA, jasmonic acid-isoleucine (JA-Ile) and abscisic acid (ABA) in rosette leaves at four, five, six, and seven weeks after sowing. We found that SA levels were significantly higher in five-week-old *mlo2* plants compared to Col-0 (1.2 vs. 0.4 nmol SA/g FW), while the *mlo3* mutant lines were mostly comparable to Col-0 (Figure 7, Figure S7, Figure S8 and Table S2). There were only a few detectable differences found in *mlo* mutant plants compared to wild type with regard to most of the jasmonates, with the exception of the JA conjugate 12-OH-JA-Ile, which is a JA-Ile degradation product and was significantly increased in a transient manner in seven-week-old *mlo3*-4 (4-fold), *mlo3*-5 (12-fold), and *mlo3*-6 (3-fold) plants. Notably, the SA conjugate SAG was in tendency increased in five-week-old *mlo3* mutant plants (*mlo3*-4 and *mlo3*-5, statistically significant with p<0.05) (Figure 7, Figure S7, Figure S8 and Table S2). We had a closer look at the SA to JA-Ile and SAG to JA-Ile ratios to assess the homeostasis of these two antagonistically acting hormones. The SAG/JA-Ile ratio was significantly increased in favor of SAG in five-week-old *mlo2*-5 (5-and 6-fold, respectively), and between 3.5 and 9-fold in *mlo3*-2, *mlo3*-4, *mlo3*-5 and *mlo3*-6 mutant plants, compared to Col-0. At six and seven weeks, only *mlo2*-5 plants displayed clearly and persistently increased SAG/JA-Ile ratios (15-fold and 10-fold, respectively). Taken together, *mlo3* mutant plants show noticeable transient differences to wild type plants in the ratio of SAG/JA-Ile in five-week-old plants.

**Fig 7.**
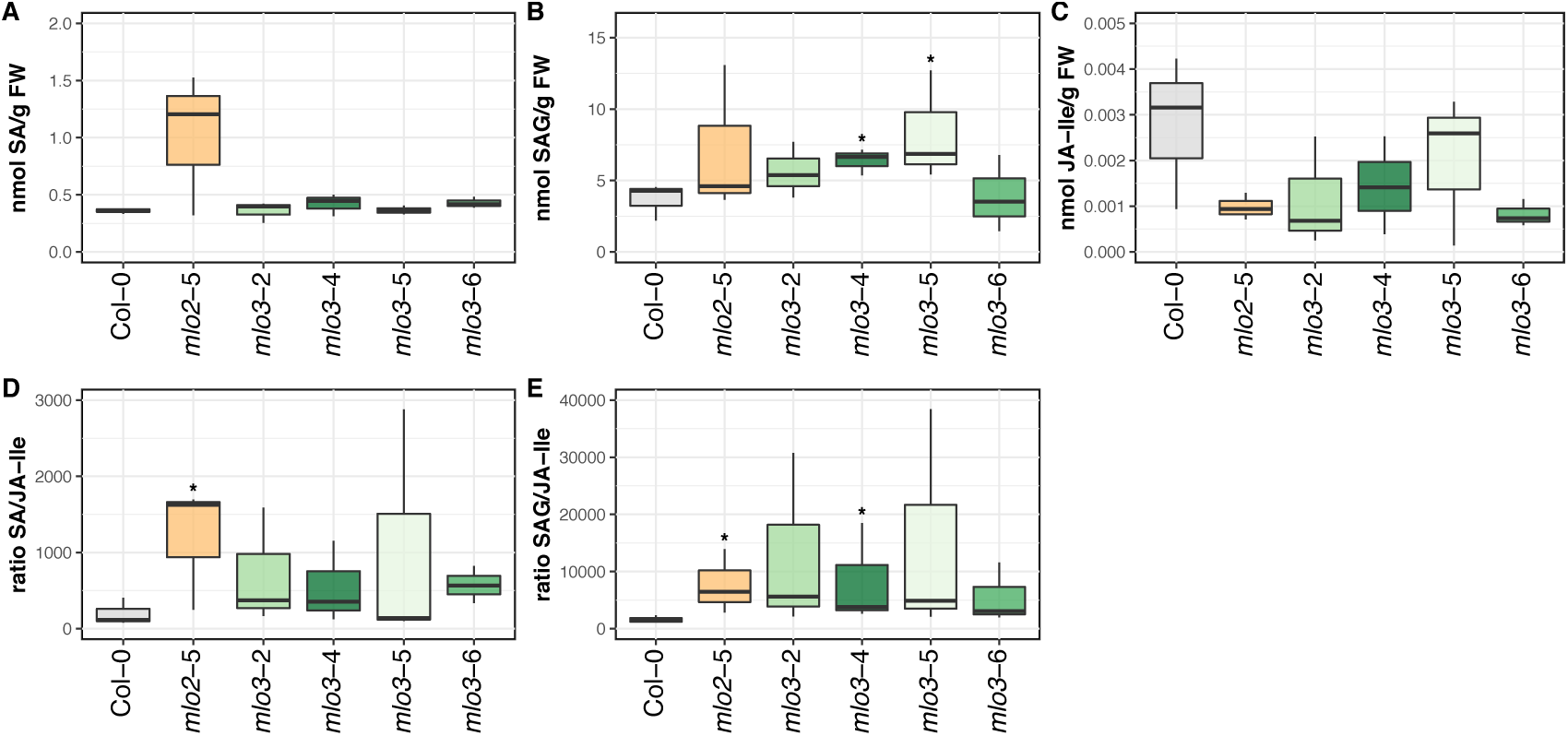
Homeostasis of the phytohormone conjugates SAG and JA-IIe are mildly affected in five-week-old *mlo3* mutant plants. The levels of the phytohormones (**A**) SA, (**B**) SA glucoside (SAG), and (**C**) JA-isoleucine (JAIle) in the leaves of unchallenged Col-0, *mlo2*-5, *mlo3*-2, *mlo3*-4, *mlo3*-5, and *mlo3*-6 plants grown under a short day cycle were measured at four, five, six, and seven weeks post germination by UPLC-nano ESI-MS/MS. Only values for five-week-old plants are shown. Based on these values, the ratio of SA over JA-Ile (**D**) and SAG over JA-Ile (**E**) were calculated for each plant. The boxplots are based on n = 3 independent replicates; statistical analysis was performed using GLM (quasi-Poisson distribution) and Kruskal rank sum tests.

## Conclusions

The functional role of most MLO family members remains enigmatic to date (20). Here we conducted a detailed characterization of the *AtMLO3* expression pattern and tested a potential contribution of the gene to plant immunity or premature senescence. Based on comprehensive *in silico* analysis (public microarray data) we found that despite some overlap *AtMLO3* is in part expressed in different tissues and under different conditions than *AtMLO2* (Figure 1A). Experimental results obtained with a P*AtMLO3*::*GUS* reporter line were, however, largely incongruent with the public microarray data, especially regarding the elevated expression levels in senescent leaves (Figure S2). This discrepancy could be due to the fact that essential *cis*-regulatory sequences may reside e.g. in *AtMLO3* up-or downstream regions and/or intronic sequences not covered by the transgenic P*AtMLO3*::*GUS* construct. It is known that in particular promoter-proximal introns in Arabidopsis genes are enriched in sequence motifs that enhance gene expression (57). These could also be critical for the native expression pattern of *AtMLO3*, in particular the senescence-associated expression, which was essentially lacking in the P*AtMLO3*::*GUS* transgenic line. The generally weak GUS staining obtained with the P*AtMLO3*::*GUS* transgenic line could also simply reflect the natural (*ca.* ten-fold) differences in *AtMLO2* and *AtMLO3* expression levels (Table S1).

Co-expression networks can be exploited to predict gene functions (58). *AtMLO2* and *AtMLO3* share a considerable number of co-expressed genes (79 out of the top 300; Figure 1B). More than 30 % of these co-expressed genes are functionally related to plant defense and/or SA signaling (Figure 1C). A set of genes co-expressed with *AtMLO2* was previously found to represent an evolutionarily conserved regulon with a function in antifungal plant innate immunity (59). Most of the co-expressed genes share a common *cis*-regulatory element in their promoters, and several mutants of these genes exhibit altered pathogen infection phenotypes and perturbed levels of antifungal defense compounds (glucosinolates and glucosinolate metabolites). Since key components of this regulon with well-defined functions in exocytosis (e.g. *AtSYP121*/*AtPEN1, AtSNAP33* and *AtVAMP722*) and glucosinolate biosynthesis/secretion (e.g. *AtPEN2, AtPDR8*/*AtPEN3, AtCYP83B1*) are lacking in the list of genes jointly co-expressed with *AtMLO2* and *AtMLO3*, the here reported 79 genes may define a separate regulon with a distinctive function. Given the overrepresentation of SA-related GO terms in the 79 genes (Figure 1C), this function could be related to SA-dependent defense and/or other SA-dependent processes. Many of the putative *cis*-regulatory elements we identified in the upstream sequences of these genes may be binding sites for WRKY transcriptional regulators. For instance, we found potential *At*WRKY18 and *At*WRKY38 binding sites to be abundant (Supplementary File 3). Notably, *wrky18 wrky40* double mutants were shown to exhibit increased resistance to *G. orontii* accompanied by de-regulated levels of SA and JA (60), while *At*WRKY38 and *At*WRKY62 both function as negative regulators of SA-dependent defense against *P. syringae* (61). These findings further support a potential role for *AtMLO3* in SA-dependent defense-related processes. Together with the strong transcriptional up-regulation of *AtMLO3* in systemic tissues following local bacterial challenge and the *AtFMO1*-dependent induction of *AtMLO3* expression upon treatment with the SAR-like and defense priming inducer pipecolic acid (Figure S1), this may point to a putative role for *AtMLO3* in SAR.

Based on the close phylogenetic relationship of *AtMLO3* and the characteristics of its expression profile outlined above, we hypothesized that *mlo3* mutants might show altered pathogen infection phenotypes. We selected four independent *mlo3* T-DNA insertion mutants that each represent a null mutant (full-length transcript lacking; Figure 2) and subjected these mutants to inoculation with various phytopathogens from three kingdoms of life. These assays revealed essentially unaltered infection phenotypes upon challenge with either adapted or non-adapted powdery mildew fungi (Figure 3), the oomycete *H. arabidopsidis* (Figure 5A) and the bacterial pathogen *P. syringae* (Figure 5B and C). Notably, in contrast to *mlo2* (11), even the SA-dependent SAR response to *P. syringae* was unaffected in *mlo3* mutants (Figure 5D). Absence of a detectable pathogen infection phenotype in our set of *mlo3* mutants might be explained in different ways. First, we only used biotrophic (powdery mildews, *H. arabidopsidis*) or hemibiotrophic (*P. syringae*) pathogens in our bioassays. It would be interesting to expand these experiments also to necrotrophic pathogens (e.g. *Botrytis cinerea*) in the future. Second, we only scored particular parameters (e.g. pathogen entry and sporulation) of each plant-pathogen interaction but did not assess others (e.g. the rate of hyphal expansion) and thus might have missed a respective (subtle) phenotype. Third, *mlo3* mutants could be affected for a given parameter in a particular plant-pathogen interaction, but the effect might be too small to be recognized with the limited number of experimental replicates we performed. Finally, it of course also remains a possibility that *mlo3* mutants do not show any altered infection phenotypes at all.

Apart from not presenting a distinctive alteration on their own, *mlo3* mutants failed to exert a synergistic (enhancing resistance) or epistatic (relieving resistance) effect on the infection phenotype of partially (*mlo2* and *mlo2 mlo6*) or fully (*mlo2 mlo6 mlo12*) powdery mildew-resistant mutant plants in any of the pathogen assays (Figure 3 and Figure S3). Unlike *AtMLO6* and *AtMLO12* (4) and despite the set of co-expressed genes shared with *AtMLO2* (see above), *AtMLO3* therefore does not seem to exhibit any cryptic and/or unequal genetic redundancy with *AtMLO2* regarding plant immunity. The absence of any pathogen phenotype in *mlo3* mutants might be considered rather unexpected given the plant immunity-related expression profile and co-expression network of the *AtMLO3* gene. However, it is consistent with the fact that so far all *mlo* mutants associated with altered pathogen defense have been ascribed to members of the phylogenetic clades IV (monocotyledonous plants) and V (dicotyledonous plants) of the MLO family, while *At*MLO3 is a member of clade VI (3, 19).

In unchallenged leaf epidermal cells, the barley Mlo protein localizes evenly to the cell periphery (plasma membrane) but accumulates in discrete areas upon attempted attack by the powdery mildew pathogen (62). Such pathogen-triggered redistribution and focal accumulation at fungal attack sites has been seen in the case of several but not all tested plasma membrane-resident proteins (62). For Arabidopsis *At*PEN1 and *At*PEN3 it was shown that both actin-dependent and –independent processes contribute to the recruitment of these proteins to regions of biotic stress (63). The focal accumulation of proteins at these sites correlates with their extracellular incorporation into local pathogen-induced cell wall compartments (papillae; (64)). While the *At*MLO2 protein, reminiscent of barley Mlo, shows focal accumulation at fungal pathogen attack sites, the *At*MLO3 protein lacks this dynamic behavior and remains evenly distributed at the cell periphery following powdery mildew challenge (Figure 4). This result suggests that the capacity for pathogen-triggered focal accumulation is not an intrinsic feature of all MLO proteins but an isoform-specific attribute. Specific peptide motifs in the protein, certain post-translational modifications and/or association with particular interaction partners may determine the membrane dynamics of individual MLO paralogs.

Powdery mildew-resistant barley *mlo* mutants are well known for their premature senescence phenotype, which is associated with spontaneous callose deposition and catabolism of photosynthetic pigments (5, 6, 8). Arabidopsis *mlo2* mutants (4, 7) and wheat *mlo* null mutants (65) share this undesired feature. Results of genetic analyses revealed that at least in Arabidopsis this phenotype is fully dependent on SA biosynthesis and signaling, as *mlo2* double mutants with *sid2, pad4* or *npr1* result in restored wild type-like leaf senescence (4). Given the SA-related co-expression profile of *AtMLO3* (Figure 1C) we explored whether *mlo3* T-DNA mutants show likewise any signs of early leaf senescence. We found that *mlo3* mutants, reminiscent of *mlo2* mutants, exhibit both premature leaf chlorosis/necrosis (Figure 6C-D) and spontaneous callose deposition in the absence of any pathogen (Figure 6A-B). The extent of the latter phenotype is, however, less pronounced than in the *mlo2*-5 null mutant and thus quantitatively intermediate between the Col-0 wild type and the *mlo2*-5 genotype (Figure 6A). Similar to powdery mildew infection (Figure 3), the absence of *AtMLO3* did not result in any detectable synergistic or epistatic effect regarding spontaneous callose deposition of the *mlo2* mutant (Figure 6B and Table 2). However, putative synergism in *mlo2 mlo3* double mutants could be masked by the comparatively small contribution of the *mlo3* mutation to the overall phenotype. By contrast, we noted that additional mutations in *AtMLO6* and *AtMLO12* appear to raise the number of callose deposits in the background of the *mlo2*-5 mutant, pointing to genetic redundancy of *AtMLO2, AtMLO6* and *AtMLO12* regarding this phenotype. In case of *mlo2* mutants, the callose synthase *At*GSL5/*At*PMR4 is required for the formation of callose deposits – *mlo2 pmr4* double mutants lack this feature (7). We demonstrate by *mlo3 pmr4* double mutant analysis that analogously *At*GSL5/*At*PMR4 is also required for spontaneous callose deposition in *mlo3* mutant plants (Figure 6B and Table 2). This suggests a common signaling pathway leading to aberrant callose deposits in the absence of *AtMLO2* or *AtMLO3*. Our findings further support the previous notion that spontaneous callose deposition is not causally linked to the powdery mildew resistance trait. In the case of the *mlo2* mutant, both aspects can be genetically uncoupled, as for example demonstrated by *mlo2 sid2, mlo2 pmr4* and *mlo2 rcd1* double mutants, which all exhibit resistance but lack the callose phenotype (4, 12). Conversely, *mlo3* mutants shows spontaneous callose deposition but lack enhanced powdery mildew resistance.

In Arabidopsis, a developmentally controlled increase in SA and SAG levels in the rosette leaves of *mlo2* and *mlo2 mlo6 mlo12* mutant plants precede the premature senescence phenotype (4, 56). As previously demonstrated by genetic analysis, SA/SAG accumulation is required for the occurrence of both, signs of leaf senescence and spontaneous callose deposition in *mlo2* mutants (4). We thus determined the levels of various phytohormones, including SA/SAG and the antagonistically acting JA and some of its derivatives, in the course of plant development. Results of this experiment confirmed the previously reported (56) developmentally controlled increase in SA/SAG levels in the *mlo2* mutant in comparison to the Col-0 wild type (Figure 7A and B). We also found transiently (in five-week-old plants) increased SAG levels and concomitantly reduced JA-Ile levels in *mlo3* mutant plants, resulting in a markedly increased SAG/JA-Ile ratio compared to Col-0 wild type (Figure 7E). Unlike the *mlo2*-5 mutant, which apart from an increased SAG/JA-Ile also showed an enhanced SA/JA-Ile ratio, the SA to JA-Ile relation was essentially wild type-like in the *mlo3* mutants (Figure 7D). It thus seems as if the *mlo3* mutants represent a weakened form of the *mlo2* mutant regarding alterations in leaf phytohormone content, with the conjugated forms (SAG and JA-Ile), but not with the unconjugated SA showing aberrant accumulation during plant development. Notably, in contrast to the *mlo2* mutant, this effect is transient in the *mlo3* mutant plants, which likely explains the much less pronounced callose phenotype seen in these genotypes (Figure 6A-B).

In summary, our study revealed dissimilar roles for *AtMLO2* and *AtMLO3* in pathogen defense but functional overlap between the two genes regarding the control of timely leaf senescence. Both genes/proteins might act cooperatively in a shared pathway involving SA signaling. Absence of either gene results in spontaneous *At*GSL5/*At*PMR4-dependent callose deposition and premature leaf chlorosis and necrosis. It thus seems as if the clade VI protein *At*MLO3 in part functionally diverged from the clade V protein *At*MLO2 after phylogenetic separation, yet both retained a shared role in development. It remains, however, a formal possibility that *mlo3* mutants are affected in agerelated resistance (ARR). ARR describes the phenomenon of enhanced immunity in mature plants as compared to young plants (66). It is effective against pathogens from different kingdoms of life and was shown to depend on SA accumulation (67, 68). Given their premature senescence, the co-expression of genes associated with SA-related processes and the altered phytohormone balance, *mlo3* and *mlo2* mutants might show aberrant ARR – a possibility that needs to be tested experimentally in the future. Since preliminary evidence indicates that MLO proteins may have the capacity to form hetero-oligomeric complexes (69), it also remains to be seen whether *At*MLO2 and *At*MLO3 operate separately in distinct protein assemblies or at least in part act cooperatively in a common complex.

## Material and Methods

### *In silico* expression analysis

Arabidopsis gene expression data for *AtMLO1* (At4g02600), *AtMLO2* (At1g11310), *AtMLO3* (At3g45290), *AtMLO6* (At1g61560), and *AtMLO12* (At2g39200) was extracted from the eFP browser (http://bar.utoronto.ca/efp/cgi-bin/efpWeb.cgi; (27)) in January 2019. Co-expression network analysis was done using ATTED-II release 2017.12.14 (http://atted.jp; (28)). GO terms were extracted with PANTHER v10, release 2018_04 (30) and PLAZA v4.0 (29) was used for GO term enrichment analysis, using false discovery rate (FDR) at p<0.05 and Fisher’s exact test. We used AthaMap (43, 44) and MEME suite v5.0.4 (45) for the detection of putative *cis*-regulatory elements.

### Plant materials

We used four independent T-DNA insertion lines for *AtMLO3* in the background of the accession Col-0, i.e. *mlo3*-2 (SALK_063243; (4)), *mlo3*-4 (SALK_027770), *mlo3*-5 (GK_787G01), and *mlo3*-6 (SALK_116848). The novel lines were acquired from the Nottingham Arabidopsis Seed Centre (NASC) and screened for presence of T-DNA according to (70). We further included the wild type accessions Columbia (Col-0) and Landsberg *erecta* (L-*er*), and the following mutants (all in Col-0 background): *mlo1*-6, *mlo2*-5 and *mlo2*-5 *mlo6*-2 *mlo12*-1 (4), *pen2*-1 *pad4*-1 *sag101*-1 (46), *eds1*-2 (48). Combinations of *mlo3*-2 and *mlo3*-4 with *mlo2*-5, *mlo6*-2, or *mlo12*-1 were generated by crossing of the respective T-DNA insertion lines. Both *mlo3*-2 and *mlo3*-4 were also combined with *pmr4*-1 (55). The previously described transgenic promoter-GUS lines P*AtMLO2*::*GUS* and P*AtMLO3*::*GUS* (22) were used for the GUS assays.

### Plant growth conditions

Arabidopsis plants were grown on SoMi513 soil (HAWITA, Vechta, Germany) in 9×9 cm pots or round pots (r = 3 cm) under a short day cycle (10 h light period set to 22 °C, 14 h darkness at 20 °C, at 80-90 % relative humidity (RH) and a light intensity of 80-100 µmol s^-1^m^-2^) until transferring them to the respective plant pathogen growth chamber. Conditions in the *H. arabidopsidis* chamber were 16 °C, 60 65 % RH and a light intensity of 100 µmol s^-1^m^-2^set to a short day cycle (8 h light, 16 h dark period). *G. orontii*-infected plants were grown under a short day cycle at 10 h light period and 14 h darkness at 20 °C, *ca.* 65 % RH and a light intensity of 100 µmol s^-1^m^-2^. For *E. pisi* infection assays, conditions were a long day cycle 12 h light and 12 h dark period at 21 °C with the light intensity at 100 µmol s^-1^m^-2^and *ca.* 60 % RH. For SAR assays, plants were grown in individual pots (5 x 5 x 6 cm) filled with a mixture of soil (Einheitserde CL ED73), vermiculite, and sand (8:1:1) and cultivated in a separate chamber in a short day cycle (10 h light at 80-100 µmol s^-1^m^-2^/14 h dark) at 20 °C and 60-68 % RH. For flowering/seed production and for GUS staining assays (see below), Arabidopsis plants were transferred to a long day chamber six weeks after germination; long day conditions were 16 light at 23 °C, 8 h darkness at 20 °C, with 60-65 % RH and the light intensity was at 105-120 µmol s^-1^m^-2^. Plants for senescence assays were exclusively grown under long day conditions.

### Gene expression analysis by RT-PCR

Leaves were collected from five-week-old Arabidopsis plants grown under a short day cycle, flash-frozen in liquid N_2_ and milled to fine powder at 30 s^-1^for 30 s. RNA isolation was performed using the NucleoSpin® RNA Plant kit (Macherey-Nagel, Düren, Germany) following the manufacturer’s instructions, and RNA integrity was determined by agarose gel electrophoresis. cDNA synthesis was done with 500 ng RNA using the High-Capacity RNA-to-cDNA kit (Applied Biosystems/Thermo Scientific). The RT-PCR was performed with *Taq* polymerase at an annealing temperature of 54 °C, an elongation time of 60 s (for *AtMLO3*) or 90 s (for *AtMLO2*), and 35 cycles. Primers for *AtMLO3* RT-PCR were AtMLO_3C (forward primer), 5’-GAAAATGGAATTCGTGGGAGAAAGAG-3’ and AtMLO_331 (reverse primer), 5’-GCTAATACAAGTCGTGTGATTATG-3’, resulting in a 697 bp product. Primers for AtMLO2 RT-PCR were AtMLO_2C, 5’-ATGGAGATGGAGATAAACCCGGTC-3’ (forward primer) and AtMLO_2bw1, 5’-ACTAGTATCTAGGAGAAGGAG-3’ (reverse primer), generating a product of 1,209 bp.

### GUS staining

The plant tissue was submerged in GUS staining solution (39 mM NaH_2_PO_4_, 61 mM Na_2_HPO_4_, 2 mM K_3_Fe(CN)_6_, 0.5 mg mL^-1^ X-Gluc (Apollo Scientific, Whitefield, UK), 0.1 % v/v Triton X-100) and vacuum was applied three times for 10 min. Then, seedlings were incubated in the solution for 2-3 h, while plant rosettes were incubated overnight (up to 16 h), both at 37 °C. Subsequently, the samples were bleached in 80 % ethanol at room temperature for at least two to three days. Photographs of GUS-stained seedlings or of GUS-stained leaves after pathogen infection were taken using the Keyence Biorevo BZ 9000 microscope (Keyence Corporation, Osaka, Japan), rosettes were photographed with a Coolpix P600 camera (Nikon, Amsterdam, The Netherlands).

### Callose staining

Arabidopsis plants were grown under a short day cycle for six weeks. Then, all leaves from the rosette were collected in 80 % ethanol for bleaching of the chlorophyll for at least three days at room temperature. Thereafter, the leaves were submerged in aniline blue staining solution (150 mM K_2_HPO_4_, 0.01 % w/v aniline blue; Sigma-Aldrich, Munich, Germany) in the dark overnight. For microscopic assessment, the leaves were first mounted on a glass slide. The samples were analyzed by fluorescence microscopy with a Keyence Biorevo BZ 9000 microscope (Keyence Corporation), using UV excitation (327-427 nm) and the DAPI/Aniline blue emission filter (emission 417-477 nm). Pictures were taken with the BZII-viewer software, set to long exposure time (1/7 to 1/3 s) and high black balance to remove background fluorescence. Callose deposit counts were performed using CellProfiler v3.1.8 software for batch analysis (53).

### Powdery mildew infection assays

Arabidopsis plants were inoculated with powdery mildew at *ca.* r = 2.5 cm in rosette size at four to five weeks after g ermination. *Golovinomyces orontii* is adapted to infection of Arabidopsis and was cultivated on susceptible *eds1*-2 plants; *Erysiphe pisi* was cultivated on susceptible pea isolates. Inoculation was conducted by leaf-to-leaf transfer of conidiospores. Leaves from at least four individual plants were collected for each replicate at 48 hpi, and five independent replicates were performed for each assay. The leaves were bleached in 80 % ethanol at room temperature for at least two to three days. Prior to microscopic analysis, fungal structures were stained by submerging the leaves in Coomassie staining solution (45 % v/v methanol, 10 % v/v acetic acid, 0.05 % w/v Coomassie blue R-250; Carl Roth, Karlsruhe, Germany) twice for 15-30 s and shortly washed in tap water in between and thereafter. The samples were analyzed with an Axiophot microscope (Carl Zeiss AG, Jena, Germany). The fungal penetration rate was determined as the percentage of spores successfully developing secondary hyphae over all spores that attempted penetration, visible by an appressorium. Macroscopic pictures of *G. orontii*-infected plants were taken at 7 dpi with a Coolpix P600 camera (Nikon).

### *Hyaloperonospora arabidopsidis* infection assays

Arabidopsis seedlings were grown on a *ca*. 10:1 soil-sand mix sterilized by microwaving for 15 min. The seedlings were grown in a thin lawn, and three pots per genotype (r = 3 cm) were used for each replicate. The seeds were stratified at 4 °C in the dark for two days and then grown under a long day cycle, covered by a lid for eight days. The plants were inoculated with *Hpa* Noco2 by spraying a spore suspension (4 x 10^4^spores mL^-1^) with four to five spray boosts onto each pot twice with a pause of 20-30 min in between. Then, the plants were incubated under a short day cycle for seven days. Thereafter, all plants from each pot were collected to determine the fresh weight with a fine balance. Subsequently, the spores were washed off with 1-2 mL of distilled water. The spore concentration was calculated using a hemocytometer by averaging the counts from ten of the 1 mm^2^ squares. The infection success was expressed as spores per gram fresh weight, determined according to the following formula:

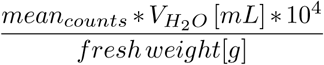

### *Pseudomonas syringae* infection assays

Arabidopsis seedlings were grown as a thin lawn, as described above for *Hpa* infection assays. The seedlings were cultivated under a short day cycle for 15 days and then placed in the dark for one day before spray-inoculation of bacteria. Prior to inoculation, the *Pst* strain DC3000 bacteria were suspended in 150 µL of 10 mM MgCl_2_, plated on NYG-Rifampicin plates, and then grown at 28 °C overnight. Afterwards, the bacteria were collected from the plates and re-suspended in 2 mL of 10 mM MgCl_2_ and diluted to an optical density at 600 nm (OD_600_) of 0.2. Directly before spray inoculation, Silwet L-77 was added to a concentration of 0.04 % v/v. Each pot was sprayed ten times with the bacteria suspension. Inoculated plants were placed in the dark for another 3-4 h before collecting 0 dpi samples. Afterwards, the plants were cultivated in a short day cycle. The bacterial titer was determined at 0 dpi as inoculation control, and at 3 dpi as follows. All plants from one pot of each genotype were collected, weighted, surface-sterilized by dipping them in 70 % ethanol, and sampled into 1.5 mL of 10 mM MgCl_2_ containing 0.01 % Silwet L-77. The bacteria were extracted by shaking at 1,500 rpm on a platform vortex for 1 h. Then, a dilution series of 10^0^ to 10^-5^ was prepared. From each dilution, 5 µL (accounting for a dilution factor of 150) were drop-inoculated on a NYG-Rifampicin plate and the plates were incubated at 28 °C for *ca.* one and a half days. The bacterial titer was expressed as log10 of colony-forming units (CFU) per gram fresh weight using the following formula:

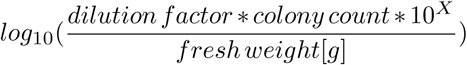

### Systemic acquired resistance assays with *Pseudomonas syringae* pv. *maculicola lux*

To monitor their SAR response, Arabidopsis *mlo3*-4, *mlo3*-6, *mlo2*-5, and Col-0 wild type plants were grown in controlled short day conditions (10 h light/14 h dark at 20 °C). The soil in which the plants were grown was microwaved for 15 min. *P. syringae* pv. *maculicola* strain ES4326 (*Psm*) and *Psm* constitutively expressing the *Photorhabdus luminescens luxCDABE* operon (*Psm lux*; (52)) were grown at 28 °C on King’s B plates containing the appropriate antibiotics. For the respective leaf inoculations, liquid King’s B cultures including antibiotics were inoculated with bacteria and grown overnight at 28 °C. After collecting the bacteria, they were washed in 10 mM MgCl_2_ three times and finally diluted to the respective optical density mentioned below. SAR experiments were conducted on five-week-old naïve plants of healthy appearance. At day one, three lower mature leaves were infiltrated from the abaxial site with either a *Psm* dilution of OD_600_ at 0.005 or 10 mM MgCl_2_ (mock control) using a needleless 1 mL syringe. Two days later, three upper mature (i.e., systemic) leaves of the pretreated plants were inoculated with a *Psm lux* dilution of OD_600_ at 0.0005 in the same way. Growth of *Psm lux* in the challenged leaves was monitored three days later using a luminometer as described in (9). Three leaves per plant were measured and after subtracting the average luminescent background, they were averaged to a value per plant expressed in relative light units (RLU) per cm^2^. The bioluminescence of *Psm lux* measured in RLUs is linearly correlated with the growth expressed in CFU (11, 52). At least seven plants per genotype and treatment were analyzed and the experiment was conducted twice.

### Transient overexpression *via* particle delivery system

Four-week-old Arabidopsis plants were transformed using the PDS-1000/He HeptaTM device (Bio-Rad, Munich, Germany) equipped with a 900-1100 psi (i.e., 6.2 to 7.6 MPa) rupture disc. Five to seven plants grown under a short day cycle covered by a lid were removed from the soil, the roots were washed with sterile water and then the plants were placed on 1 % agar containing benzimidazole. Plasmid DNA purified from a 200 mL culture using the NucleoBond® Xtra MIDI kit (Macherey-Nagel) following the manufacturer’s instructions and diluted to 1 µg µL^-1^ was used for coating of the gold particles. Then, 5 µg of plasmid DNA were bound to 3 mg gold micro-carriers (1 µm, Bio-Rad) by adding 50 µL of 2.5 M CaCl_2_ and 20 µL of 0.1 M spermidine, followed by vortexing on a platform vortex at 1,500 rpm for 3 min. The gold was pelleted by centrifugation for 2 s, first washed with 140 µL of mbox70 % ethanol and then with 140 µL of 100 % ethanol. The carriers were suspended in 60 µL of 100 % ethanol. Each of the seven macro-carriers was loaded with 7 µL of the carrier suspension. Particle delivery was carried out as recommended by the PDS-1000/He manual, applying vacuum at −24 inch Hg (−81 kPa) for Arabidopsis with the plate in the lowest level of the device. Plants were subsequently placed in the *G. orontii* infection chamber for recovery and inoculated with *G. orontii* spores one day later. Leaf samples were analyzed by confocal laser scanning microscopy at 2 dpi.

### Confocal laser scanning microscopy

Confocal laser scanning microscopy was conducted with a Leica TCS SP8 using the LAS-X software package (Leica microsystems, Wetzlar, Germany). Arabidopsis leaves were fixed on glass slides by use of double-sided tape. Fluorescence pictures and Z-stacks were recorded with the 20x or 63x water immersion objective (Leica). Fluorescence was recorded with the following specifications: YFP excitation at 514 nm (argon laser), emission at 525-570 nm.

### Osmotic stress test

Arabidopsis seeds were sterilized as follows: around 50-100 seeds were placed in a reaction tube column and submerged in 500 µL of 70 % ethanol for 1-2 min at room temperature. Next, the column was centrifuged at 1,500g for 1 min and the flow-through was discarded. The seeds were then washed with 500 µL of 100 % ethanol, centrifuged again for 1 min and after removal of the flow-through, dried by another centrifugation for 2 min. Seeds sterilized by this procedure were sown on ½ MS agar plates without (control) or containing 300 mM mannitol to apply osmotic stress. Seeds were vernalized at 4 °C in the dark overnight and then seedlings were grown under a short day cycle for three weeks, with plates being placed upright in a rack. Phenotyping was performed by measuring the root length, number of lateral roots, fresh weight, and by counting the number of leaves.

### Phytohormone determination by nanoflow Ultrahigh--Pressure Liquid Chromatography-Tandem Mass Spectrometry (UPLC-nano ESI-MS/MS)

Extraction was performed as previously described for lipids, with some modifications (71). Plant material (100 mg) was extracted with 0.75 mL of methanol containing 10 ng D4-SA, 10 ng D5-JA (both from C/D/N Isotopes Inc., Pointe-Claire, Canada), 10 ng D4-JA-Leu (kindly provided by Otto Miersch, Halle/Saale, Germany), each as internal standard. After vortexing, 2.5 mL of methyl-tert-butyl ether (MTBE) were added and the extract was shaken for 1 h at room temperature. For phase separation, 0.6 mL water were added. The mixture was incubated at room temperature for 10 min and centrifuged at 450g for 15 min. The upper phase was collected and the lower phase was re-extracted with 0.7 mL methanol/water (3:2.5, v/v) and 1.3 mL MTBE as described above. The combined upper phases were dried under streaming nitrogen and resuspended in 100 µL of acetonitrile/water (20:80, v/v) containing 0.3 mmol L^-1^ NH_4_HCOO (adjusted to pH 3.5 with formic acid).

Reversed phase separation of constituents was achieved by UPLC using an ACQUITY UPLC® system (Waters Corp., Milford, MA, USA) equipped with an ACQUITY UPLC® HSS T3 column (100 mm x 1 mm, 1.8 µm; Waters Corp., Milford, MA, USA). Aliquots of 10 µL were injected in a partial loop with needle overfill mode. Elution was adapted according to (72). Solvent A and B were water and acetonitrile/water (90:10, v/v), respectively, both containing 0.3 mmol L^-1^NH_4_HCOO (adjusted to pH 3.5 with formic acid). The flow rate was 0.16 mL min^-1^ and the separation temperature was constantly at 40 °C. Elution was performed isocratically for 0.5 min at 10 % solution B, followed by a linear increase to 40 % solution B in 1.5 min; this condition was held for 2 min, followed by a linear increase to 95 % solution B in 1 min; this condition was held for 2.5 min. The column was re-equilibrated to start conditions in 3 min.

Nanoelectrospray ionization (nanoESI) analysis was achieved using a chip ion source (TriVersa Nanomate®; Advion BioSciences, Ithaca, NY, USA). For stable nanoESI, 70 µL min^-1^ of 2-propanol/acetonitrile/water (70:20:10, v/v/v) containing 0.3 mmol L^-1^ NH_4_HCOO (adjusted to pH 3.5 with formic acid) delivered by a Pharmacia 2248 HPLC pump (GE Healthcare, Munich, Germany) were added just after the column via a mixing tee valve. By using another post column splitter 502 nL min^-1^ of the eluent were directed to the nanoESI chip with 5 µm internal diameter nozzles. Ionization voltage was set to −1.7 kV. Phytohormones were ionized in a negative mode and determined in scheduled multiple reaction monitoring mode with an AB Sciex 4000 QTRAP® tandem mass spectrometer (AB Sciex, Framingham, MA, USA). Mass transitions were as previously described (73), with some modifications and were as listed in Table S3. The mass analyzers were adjusted to a resolution of 0.7 amu full width at half-height. The ion source temperature was 40 °C, and the curtain gas was set to 10 (given in arbitrary units). Quantification was carried out using a calibration curve of intensity (m/z) ratios of [unlabeled]/[deuterium-labeled] vs. molar amounts of unlabeled (0.3-1000 pmol).

### Statistical analysis of the data

The statistics program R v.3.5.1 (R foundation, www.r-project.org) was used for statistics and generation of boxplots. Plotting was done with the ggplot function from the ggplot2 library (74). Due to the nature of our data, i.e. non-normal distribution, unequal variance, and small sample sizes of usually maximally six independent replicates, non-parametric statistical testing was devised. Statistical analyses were performed *via* Generalized Linear Modeling (GLM) assuming Poisson or binomial distribution as indicated accordingly (75), and supported by Wilcoxon-Mann-Whitney rank sum test if at least four independent replicates were available for testing. For phytohormone measurements, Kruskal rank sum tests were performed in addition.

## ACKNOWLEDGEMENTS

This study was supported by the German Research Foundation (Deutsche Forschungsgemeinschaft; DFG) [grant PA861/11-1 to R.P. and grant INST 186/822-1 to I.F.]. *H. arabidopsidis* Noco2 was kindly provided by Jane Parker (Max Planck institute for Plant Breeding Research, Cologne, Germany). The two *P. syringae* pv. *maculicola* strains were kindly provided by Jürgen Zeier (Heinrich Heine University, Düsseldorf, Germany). This work would not have been possible without coffee and chocolate.

## AUTHOR CONTRIBUTIONS

S.K. and S.T. performed the pathogen assays, K.G. did the SAR experiments. S.K. and A.R. conducted callose, senescence, GUS and osmotic stress assays. Phytohormone measurements were performed by K.Z. and K.Z. and I.F. analyzed the data. S.K. did the expression analysis, statistical testing, and plotting of the data. R.P. and S.K. designed the project and wrote the manuscript. All authors have read and approved the manuscript.

## Supplementary data

**Table S1.** *AtMLO3* is upregulated in systemic leaves of Arabidopsis accession Col-0 during SAR and in response to application of pipecolic acid (Pip). The table was compiled on the basis of data from Bernsdorff et al. (2016) and Hartmann et al. (2018). Expression is shown as reads per million (RPM), asterisks indicate significant expression differences.

| Name | AGI code | Col-0 |  |  | *fmo1*<sup>a</sup> |  |  | Col-0 |  |  |
| --- | --- | --- | --- | --- | --- | --- | --- | --- | --- | --- |
|  |  | H<sub>2</sub>O | Pip<sup>b</sup> | logFC<sup>c</sup> | H<sub>2</sub>O | Pip<sup>b</sup> | logFC<sup>c</sup> | Mock | *Psm* | logFC<sup>c</sup> |
| *AtMLO1* | At4g02600 | 24.1 | 32.2 | 0.4 | 28.0 | 29.1 | 0.1 | 32.7 | 39.3 | 0.3 |
| *AtMLO2* | At1g11310 | 55.6 | 145.1 | 1.4** | 48.3 | 52.7 | 0.1 | 78.6 | 515 | 2.7** |
| ***AtMLO3*** | At3g45290 | 5.1 | 16.5 | **1.5\*\*** | 5.2 | 5.2 | 0.0 | 10.3 | 57.5 | **2.4\*\*** |
| *AtMLO4* | At1g11000 | 6.8 | 9.3 | 0.4 | 7.7 | 8.3 | 0.1 | 6.2 | 8.2 | 0.4 |
| *AtMLO5* | At2g33670 | 0.0 | 0.0 | 0.0 | 0.0 | 0.0 | 0.0 | 0.0 | 0.0 | 0.0 |
| *AtMLO6* | At1g61560 | 0.9 | 2.8 | 1.0 | 0.8 | 0.7 | 0.0 | 1.1 | 19.2 | 3.3** |
| *AtMLO7* | At2g17430 | 1.3 | 2.4 | 0.5 | 1.3 | 1.7 | 0.2 | 1.2 | 2.1 | 0.5 |
| *AtMLO8* | At2g17480 | 13.3 | 14.8 | 0.1 | 13.4 | 15.6 | 0.2 | 20.4 | 20.2 | 0.0 |
| *AtMLO9* | At1g42560 | 0.1 | 0.1 | 0.0 | 0.2 | 0.2 | -0.1 | 0.0 | 0.0 | 0.0 |
| *AtMLO10* | At5g65970 | 4.0 | 5.2 | 0.3 | 3.3 | 3.7 | 0.1 | 2.9 | 4.0 | 0.4 |
| *AtMLO11* | At5g53760 | 15.0 | 13.5 | -0.1 | 15.2 | 15.9 | 0.1 | 14.2 | 6.7 | -1.0* |
| *AtMLO12* | At2g39200 | 0.3 | 0.4 | 0.1 | 0.5 | 0.3 | -0.2 | 0.6 | 5.7 | 2.1** |
| *AtMLO13* | At4g24250 | 0.9 | 1.4 | 0.4 | 0.8 | 1.1 | 0.2 | 0.5 | 0.5 | 0.0 |
| *AtMLO14* | At1g26700 | 0.0 | 0.0 | 0.0 | 0.0 | 0.0 | 0.0 | 0.0 | 0.0 | 0.0 |
| *AtMLO15* | At2g44110 | 0.0 | 0.0 | 0.0 | 0.0 | 0.0 | 0.0 | 0.0 | 0.0 | 0.0 |

<sup>a</sup> *fmo1*, Arabidopsis mutant of *FLAVIN MONOOXYGENASE1* (*FMO1*)
<sup>b</sup> Pip, Pipecolic acid applied by watering
<sup>c</sup> logFC, log<sub>2</sub> fold change

**Table S2.**
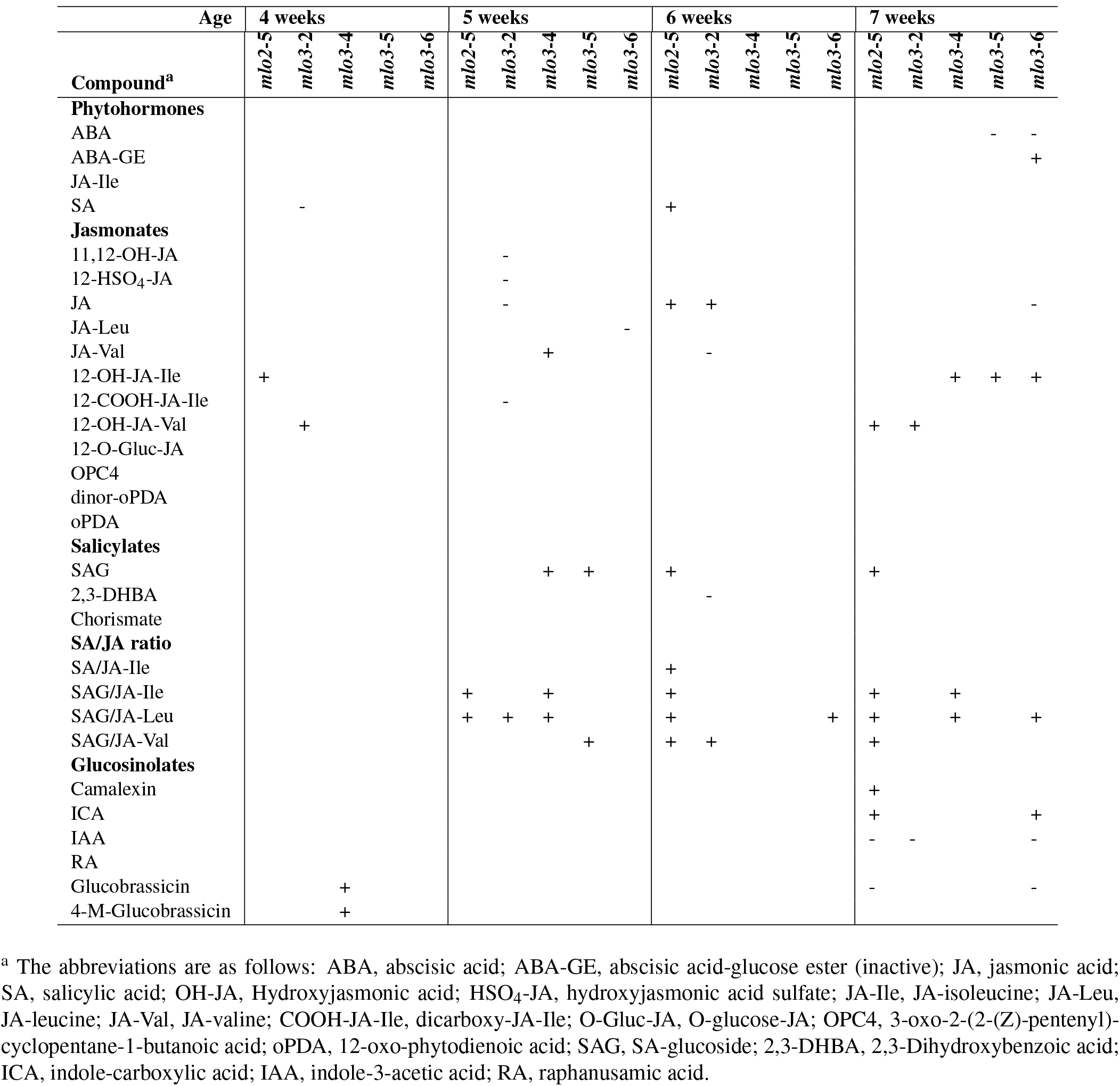
Summary of significant differences in phytohormone compound compositions in *mlo2* and *mlo3* compared to Col-0.

**Table S3.** Mass transitions and corresponding conditions for determination of the phytohormones.

| MRM transitions |  |  |  |  |  |
| --- | --- | --- | --- | --- | --- |
| Q1<sup>a</sup> | Q3<sup>a</sup> | Analyte | DP<sup>b</sup> | EP<sup>c</sup> | CE<sup>d</sup> |
| 137 | 93 | SA | -25 | -6 | -20 |
| 141 | 97 | D4-SA | -25 | -6 | -22 |
| 153 | 109 | 2,3-DHBA | -25 | -9 | -18 |
| 160 | 116 | ICA | -40 | -6.5 | -22 |
| 162 | 58 | RA | -15 | -6 | -14 |
| 174 | 130 | IAA | -35 | -9 | -14 |
| 179 | 135 | D5-IAA | -35 | -9 | -14 |
| 207 | 137 | Chorismat-H<sub>2</sub>O | -20 | -9 | -16 |
| 209 | 59 | JA | -30 | -4.5 | -24 |
| 214 | 62 | D5-JA | -35 | -8.5 | -24 |
| 225 | 59 | 11,12-OH-JA | -35 | -9 | -28 |
| 237 | 165 | OPC4 | -45 | -6 | -24 |
| 263 | 153 | ABA | -35 | -4 | -14 |
| 263 | 165 | dinor-oPDA | -40 | -5 | -20 |
| 293 | 179 | D6-ABA | -80 | -10 | -42 |
| 296 | 170.2 | D5-oPDA | -65 | -4 | -28 |
| 299 | 137 | SAG | -30 | -4 | -18 |
| 305 | 97 | 12-HSO<sub>4</sub>-JA | -30 | -4 | -32 |
| 308 | 116 | JA-Val | -45 | -5 | -28 |
| 322 | 130 | JA-Ile/Leu | -45 | -5 | -28 |
| 325 | 133 | D4-JA-Leu | -80 | -4 | -30 |
| 324 | 116 | 12OH-JA-Val | -45 | -10 | -30 |
| 338 | 130 | 12OH-JA-Ile | -45 | -10 | -30 |
| 352 | 130 | 12COOH-JA-Ile | -45 | -10 | -30 |
| 387 | 59 | 12-O-Gluc-JA | -85 | -9 | -59 |
| 425 | 263 | ABA-GE | -30 | -10 | -16 |
| 447 | 97 | Glucobrassicin | -45 | -7 | -40 |
| 477 | 97 | 4-M-glucobrassicin | -55 | -5 | -38 |

<sup>a</sup> Q1, mass of precursor ion; Q3, mass of product after fragmentation
<sup>b</sup> DP, declustering potential
<sup>c</sup> EP, entrance potential
<sup>d</sup> CE, collision energy

**Supplementary Files.** List of supplementary files.

**Supplementary File 1.** Co-expressed genes of *AtMLO1, AtMLO2*, and *AtMLO3* after ATTED-II release 2017.12.14. **Supplementary File 2.** GO terms of the 79 genes from the common co-expression network of *AtMLO2* and *AtMLO3*.

**Supplementary File 3.** *Cis*-regulatory elements of the 79 genes from the common co-expression network of *AtMLO2* and *AtMLO3* predicted by AthaMap.

**Supplementary File 4.** *Cis*-regulatory elements of the 79 genes from the common co-expression network of *AtMLO2* and *AtMLO3* predicted by MEME v5.0.4.

## Abbreviations

bp: base pair
CFU: colony forming units
dpi: days post inoculation
GO: gene ontology
GUS: β-glucuronidase
*Hpa*: *Hyaloperonospra arabidopsidis*
hpi: hours post inoculation
JA: jasmonic acid
JA-Ile: jasmonic acid isoleucine
MAMP: microbe-associated molecular pattern
MS: Murashige and Skoog
*Psm*: *Pseudomonas syringae* pv. *maculicola*
*Pst*: *Pseudomonas syringae* pv. *tomato*
RH: relative humidity
RLU: relative light units
RT-PCR: reverse transcriptase polymerase chain reaction
SA: salicylic acid
SAG: sal-icylic acid glucoside
SAR: systemic acquired resistance
T-DNA: Transfer-DNA
UPLC-nano ESI-MS/MS: Ultrahigh-Pressure Liquid Chromatography-Tandem Mass Spectrometry
YFP: yellow fluorescent protein

