## Supplementary material for "Arabidopsis *mlo3* mutant plants exhibit spontaneous callose deposition and signs of early leaf senescence": Figure S1

**Supplementary Figure 1. Developmental map of *AtMLO3* transcript abundance.** The developmental map was taken from the Arabidopsis eFP browser at http://bar.utoronto.ca/efp2/Arabidopsis/Arabidopsis_eFPBrowser2.html (Winter et al., 2007). **A.** Developmental map of *AtMLO3* expression. **B.** Developmental map of *AtMLO2* expression. The expression levels are shown as absolute expression values, as indicated by the scale. The respective tissues are indicated below their graphical representation. Data was downloaded in January 2019.
