## Supplementary material for "Arabidopsis *mlo3* mutant plants exhibit spontaneous callose deposition and signs of early leaf senescence": Figure S3

**Supplementary Figure 3. Arabidopsis *mlo3* mutants show unaltered powdery mildew symptoms.** Arabidopsis *mlo3* mutant lines and mutant combinations of *mlo3*-4 with *mlo2*-5, *mlo6*-2 and *mlo12*-1 were inoculated with *G. orontii* and photographs were taken at 7 dpi. Col‑0 is the wild type, *mlo2*‑5 the resistant control. The first row shows the single mutants, the second row controls possessing wild type *AtMLO3*, the third row the respective *mlo3*‑4 mutant combinations with *mlo2*‑5, *mlo6*‑2, and *mlo12*‑1. Scale bar: 1 cm.
