## Supplementary material for "Arabidopsis *mlo3* mutant plants exhibit spontaneous callose deposition and signs of early leaf senescence": Figure S4

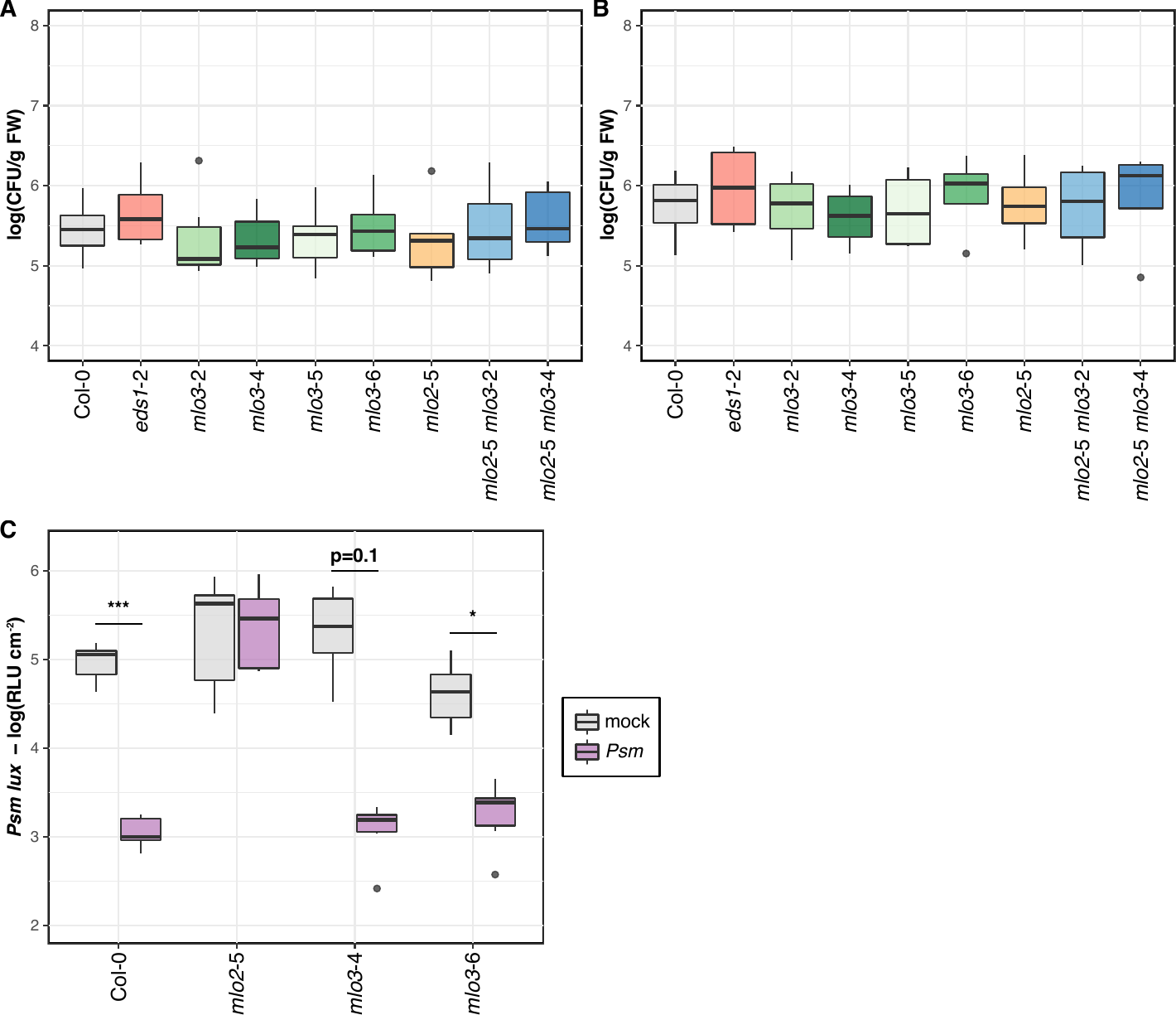

**Supplementary Figure 4. Systemic acquired resistance against *Pseudomonas syringae* is not affected in *mlo3* mutants.** For each *Pst* inoculation assay (**Fig. 5B** and **5C**), an inoculation control was measured at *ca*. 3‑4 hpi (= 0 dpi). Genotypes are as in (**Fig. 5B** and **5C**). **A.** 0 dpi control for *Pst* DC3000, n = 6 independent replicates. **B.** 0 dpi control for *Pst* DC3000 Δ*hrcC*, n = 4 independent replicates. **C.** The SAR assay was conducted the same way as in **Fig. 5D**. SAR was monitored by primary infiltrations of three lower leaves of five-week-old plants with *Psm* or 10 mM MgCl_2_ (mock), followed by inoculations of systemic leaves with *Psm lux* two days later. Bacterial proliferations of *Psm lux* were measured at 3 dpi, expressed as relative light units (RLU) cm^-2^. n = 1 biological replicate with seven plants per genotype, statistical testing was performed using GLM (quasi-Poisson distribution).
