## Supplementary material for "Arabidopsis *mlo3* mutant plants exhibit spontaneous callose deposition and signs of early leaf senescence": Figure S5

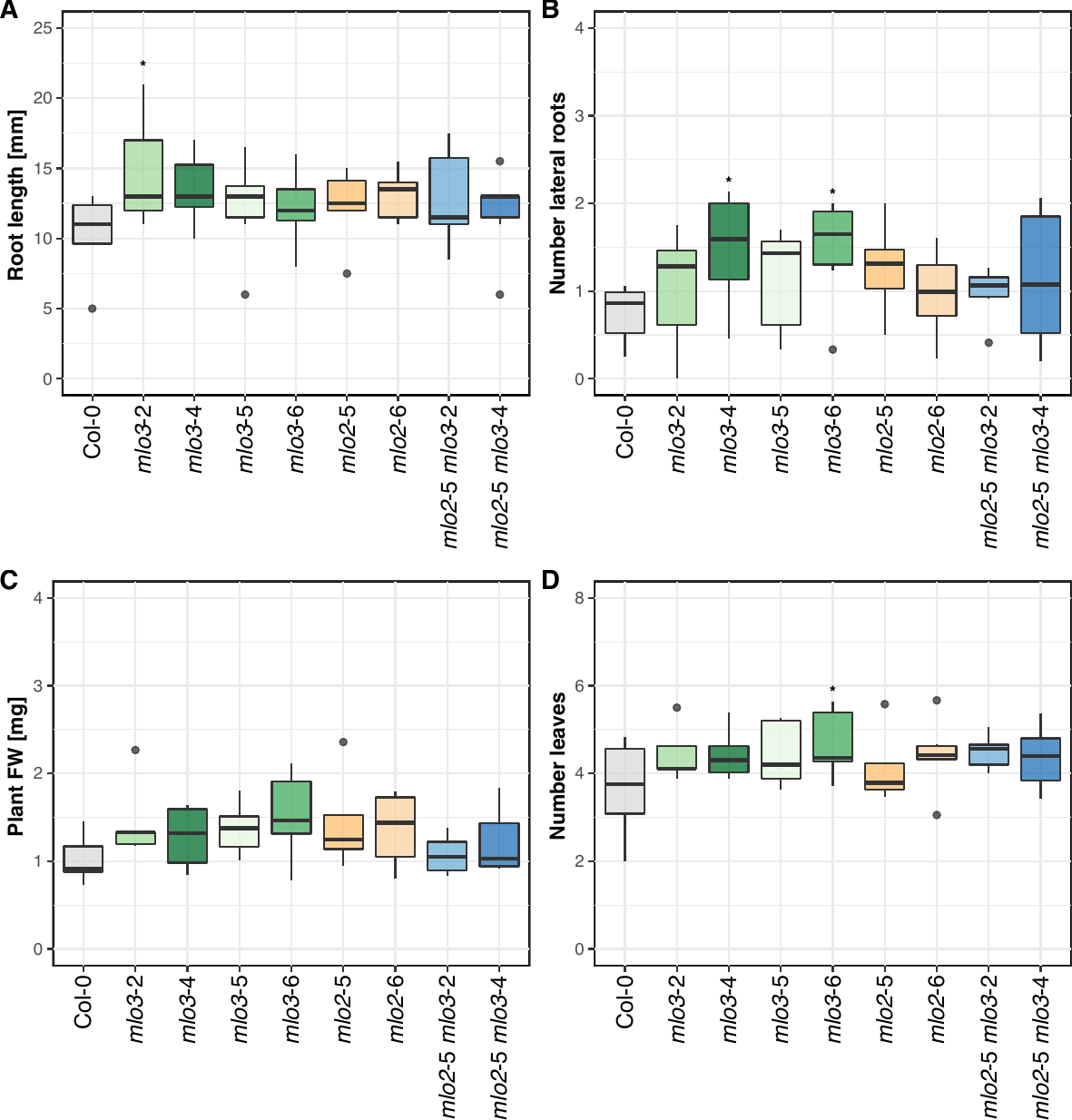

**Supplementary Figure 5. Mutants of *AtMLO3* are not altered in the osmotic stress response.** Seedlings were cultivated *in vitro* on ½ MS plates containing 300 mM of mannitol for 21 d. Genotypes were the same as in **Fig. 5B**, excluding *eds1*‑2, where Col‑0 is the wild type control, and *mlo2*‑5 is the parent control for the *mlo2 mlo3* double mutants. The following phenotypic markers were quantified: root length in mm (**A**), number of lateral roots (**B**), fresh weight in mg per plant (**C**), and number of leaves per plant (**D**). n = 5 independent replicates, statistical testing was done using GLM (Poisson distribution).
