## Supplementary material for "Arabidopsis *mlo3* mutant plants exhibit spontaneous callose deposition and signs of early leaf senescence": Figure S6

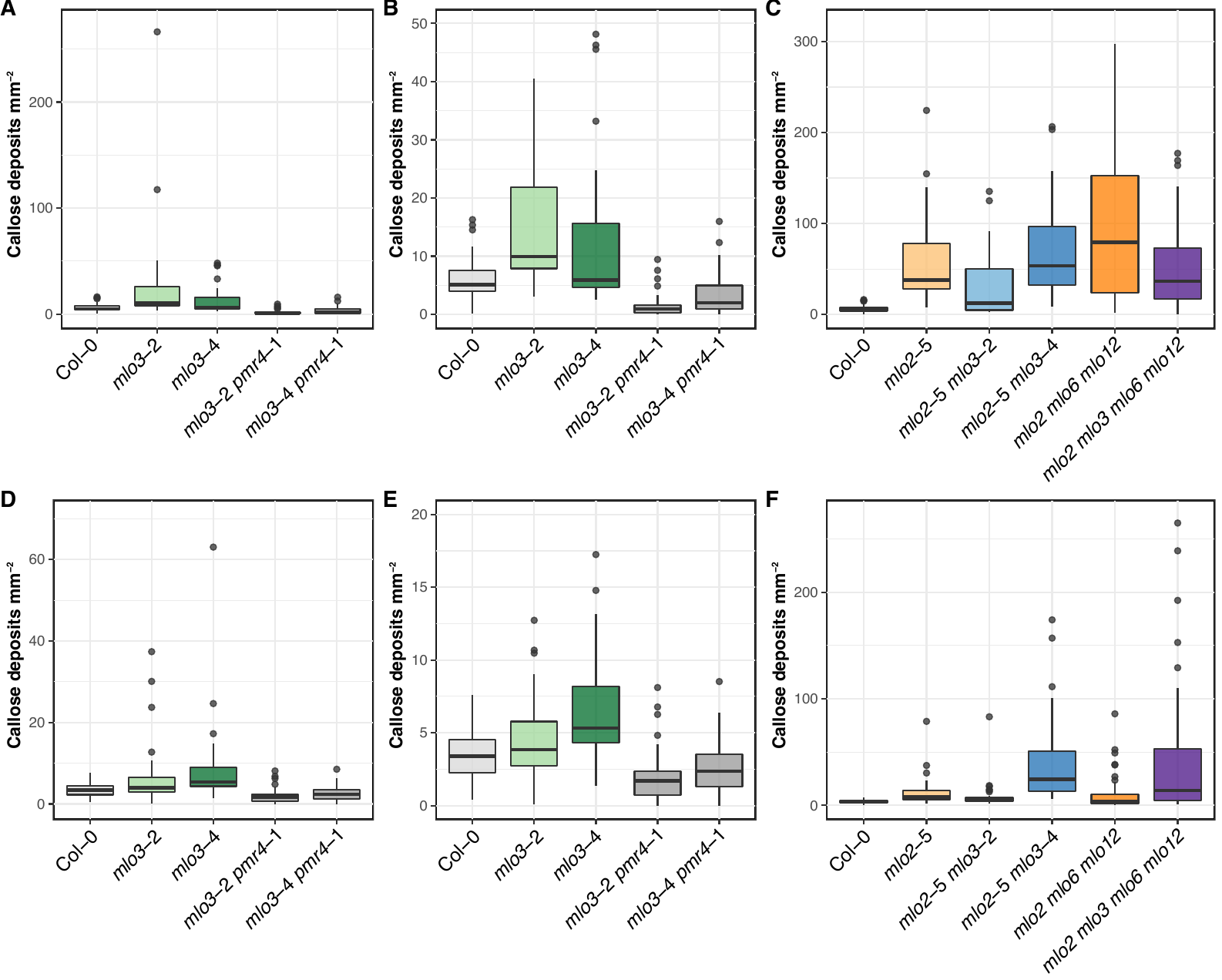

**Supplementary Figure 6. Arabidopsis *mlo3 pmr4* mutants do not show spontaneous callose deposition.** All leaves from six-week-old rosettes were collected and callose was stained using aniline blue. Col‑0 wild type served as negative control, while *mlo2*‑5 (**A**) and *mlo2*‑5 *mlo6*‑2 *mlo12*‑1 (**B**) mutants were included as positive controls. **A.** We tested *mlo3*‑2, *mlo3*‑4, and the respective double mutants with *pmr4*-1. **B.** We scaled the y-axis to highlight the differences between the genotypes. **C.** We analyzed *mlo2*‑5 *mlo3* double and the *mlo2 mlo3 mlo6 mlo12* quadruple mutants. The Col‑0 wild type control is identical in both plots. **D.**, **E.** and **F.** represent a second replicate for the experiment shown in **A**-**C**. For quantitative assessment, pictures were taken with high black balance and long exposure time to minimize background, and callose deposits were quantified using CellProfiler v3.1.8 (Carpenter et al., 2006). Each boxplot shows all counts from n = 1 independent replicate from at least 25 leaves per genotype. The mean, standard deviation and maximum value for each genotype and replicate are listed in **Table 2**.
