## Supplementary material for "Arabidopsis *mlo3* mutant plants exhibit spontaneous callose deposition and signs of early leaf senescence": Figure S7

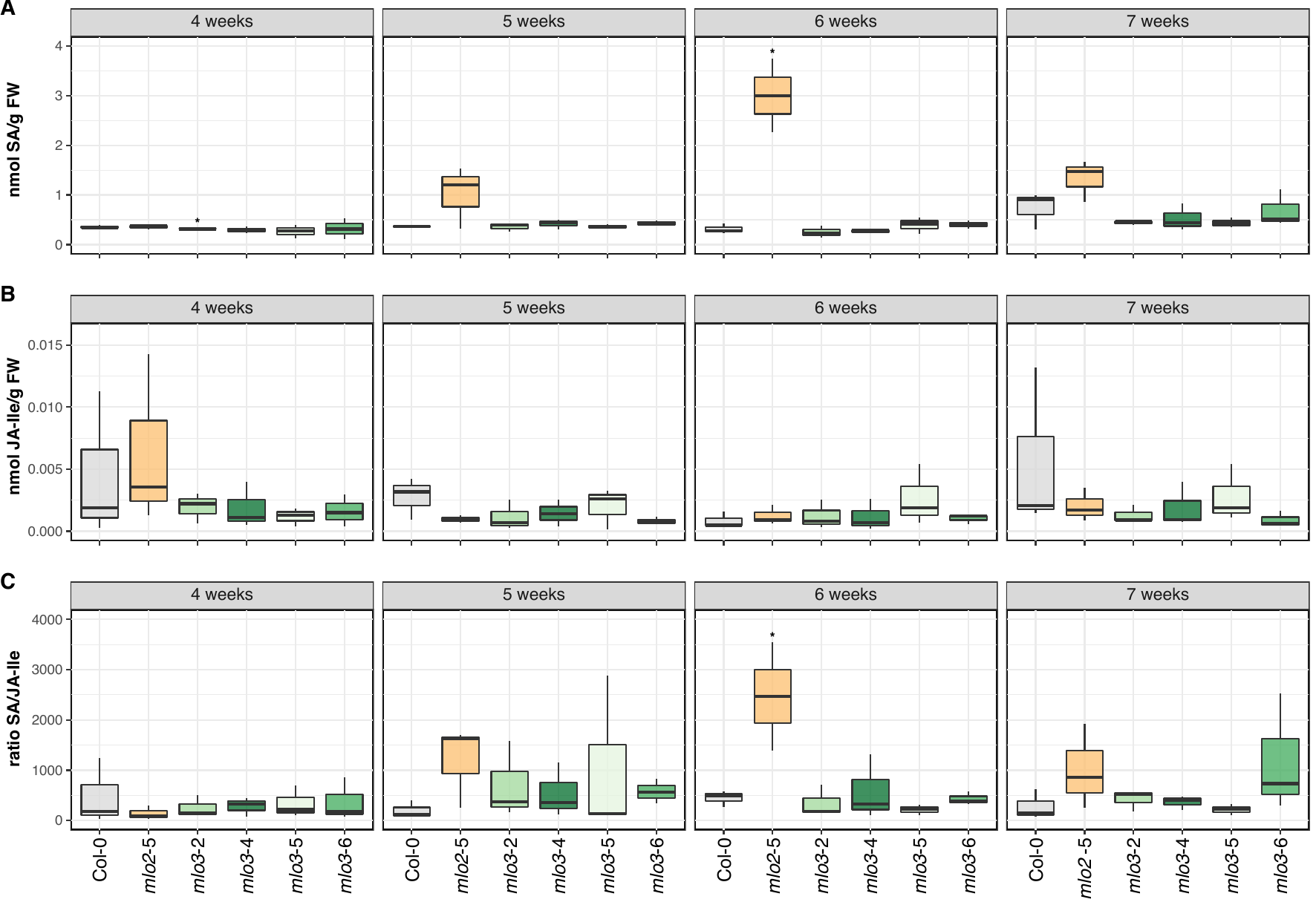

**Supplementary Figure 7. Homeostasis of SA and JA-Ile is not affected in *mlo3* mutant plants.** The level of the phytohormones (**A**) salicylic acid (SA) and (**B**) jasmoic acid (JA-Ile) in the leaves of unchallenged Col‑0, *mlo2*‑5, *mlo3*‑2, *mlo3*‑4, *mlo3*‑5, and *mlo3*‑6 plants grown under a short day cycle was measured four, five, six, and seven weeks post germination by UPLC-nano ESI-MS/MS. (**C**) Based on these values, the ratio of SA over JA-Ile was calculated for each plant. n = 3 independent replicates; statistical analysis was performed using GLM (quasi-Poisson distribution) and Kruskal rank sum tests.
