## Supplementary material for "Arabidopsis *mlo3* mutant plants exhibit spontaneous callose deposition and signs of early leaf senescence": Figure S2

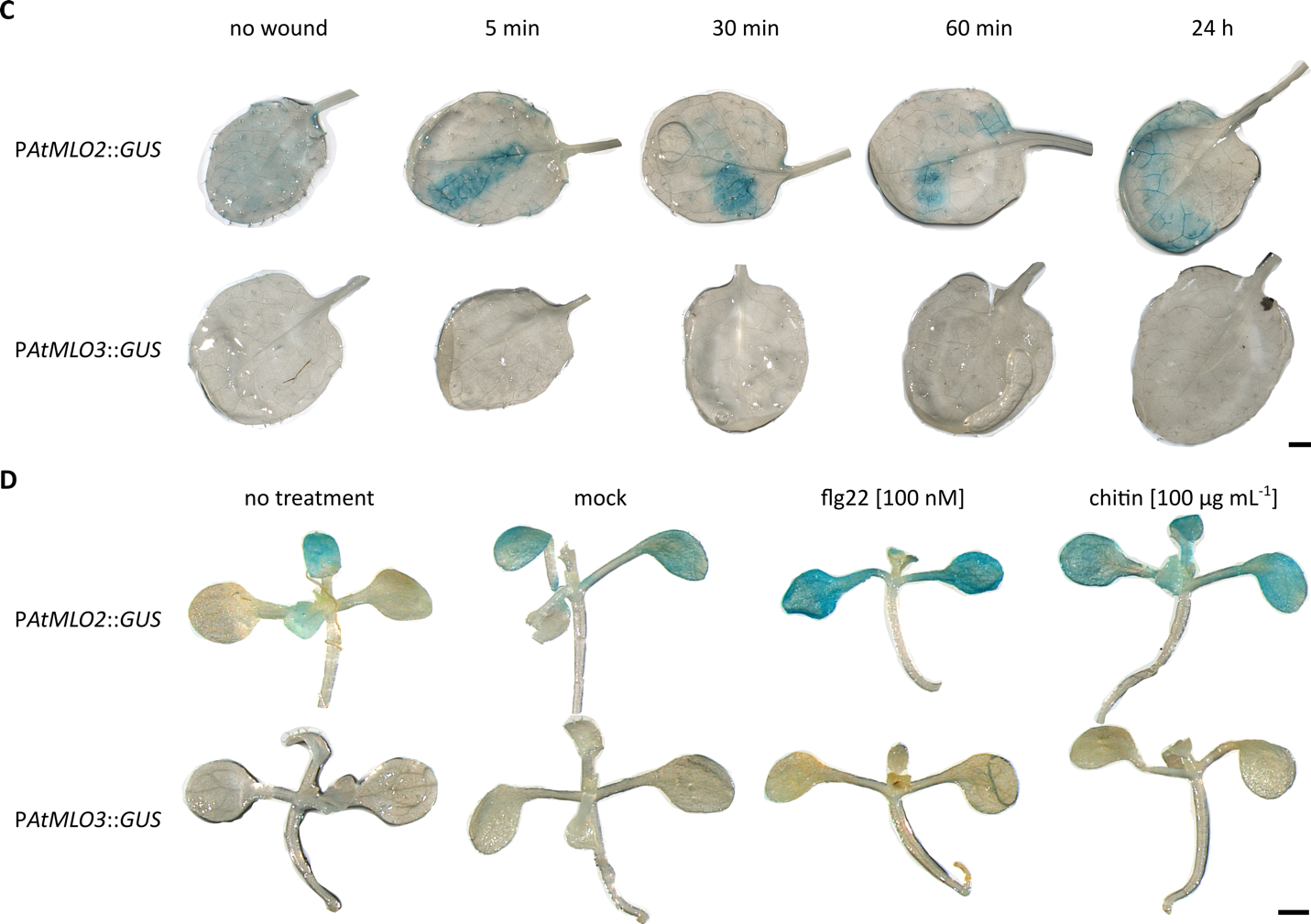

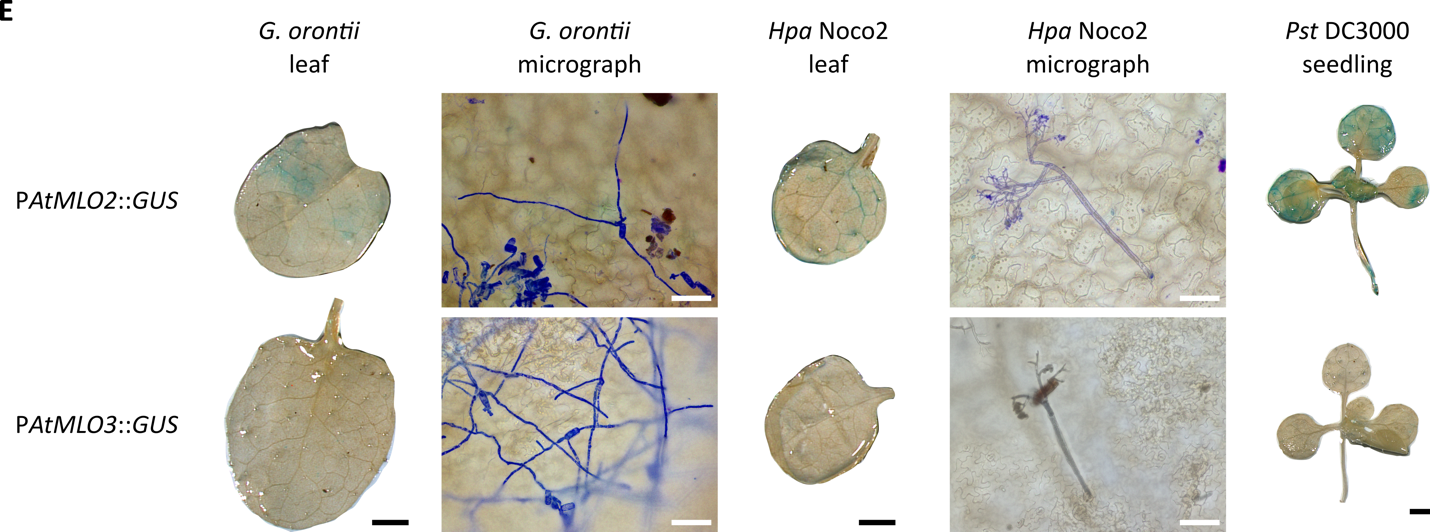

**Supplementary Figure 2. *AtMLO3* expression patterns as revealed by analysis of promoter-GUS lines.** The promoter-GUS lines P*AtMLO2*::*GUS* and P*AtMLO3*::*GUS* (Chen et al., 2006) were grown under the indicated conditions and then subjected to GUS staining and subsequent bleaching for photography. Conditions are as follows (upper panel always P*AtMLO2*::GUS and lower panel P*AtMLO3*::GUS). **A.** Rosettes from three- to eight-week-old plants, grown under a long day cycle. **B.** Rosettes from three- to eight-week-old plants, grown under a short day cycle. **C.** Leaves after wounding by tweezers. From left to right: no wounding, 5 min, 30 min, 60 min, and 24 h after wounding. **D.** Seedlings after MAMP treatment, from left to right: no treatment, mock treatment, flg22 (100 nM), chitin (100 µg mL^-1^). **E.** Leaves after pathogen infection, from left to right: *G. orontii* leaf, *G. orontii* micrograph, *Hpa* leaf, *Hpa* micrograph, *Pst* DC3000 leaf. The three- or four-week-old plants from (**A**) were the unchallenged controls here. All conditions were tested three to five times with similar results, representative pictures are shown. Scale bars of micrographs: 100 µm; scale bars of photographs: 1 cm.
